# All spectral frequencies of neural activity reveal semantic representation in the human anterior ventral temporal cortex

**DOI:** 10.1101/2025.04.17.649404

**Authors:** Saskia L. Frisby, Ajay D. Halai, Christopher R. Cox, Alex Clarke, Akihiro Shimotake, Takayuki Kikuchi, Takeharu Kuneida, Yoshiki Arakawa, Ryosuke Takahashi, Akio Ikeda, Riki Matsumoto, Timothy T. Rogers, Matthew A. Lambon Ralph

**Affiliations:** MRC Cognition and Brain Sciences Unit, University of Cambridge, Cambridge, UK; Department of Psychology, Louisiana State University, Baton Rouge, USA; Department of Psychology, University of Warwick, Warwick, UK; Department of Neurology, Kyoto University Graduate School of Medicine, Kyoto, Japan; Clinical Research Center, National Hospital Organization Utano Hospital, Kyoto, Japan; Department of Neurosurgery, Kyoto University Graduate School of Medicine, Kyoto, Japan; Department of Neurosurgery, Ehime University Graduate School of Medicine, Ehime, Japan; Department of Epilepsy, Movement Disorders and Physiology; Kyoto University Graduate School of Medicine; Kyoto; Japan; Division of Neurology, Kobe University Graduate School of Medicine, Kusunoki-cho, Kobe, Japan; Department of Psychology, University of Wisconsin–Madison, Madison, USA

**Keywords:** electrocorticography, ECoG, intracranial electrophysiology, time-frequency analysis, semantic representation, decoding, multivariate pattern analysis

## Abstract

Intracranial electrophysiology offers a unique insight into the nature of information representation in the brain – it can be used to disentangle information encoded in local neuronal activity (gamma and high gamma frequencies) from information encoded via long-range interactions (lower frequencies). We used regularised logistic regression to decode animacy from time-frequency power and phase extracted from electrocorticography (ECoG) grid electrode data recorded on the surface of human vATL. Power in gamma (30 – 60 Hz) and high gamma (60 – 200 Hz) produced reliable decoding, indicating that semantic information is indeed expressed by local populations in vATL. However, power from a wide range of frequencies (4 – 200 Hz) produced significantly higher decoding accuracy and also exhibited the same rapidly-changing dynamic code previously observed when decoding voltage. These findings support the theory that semantic information is encoded by a local vATL “hub” that interacts with distributed cortical “spokes”.

---

Semantic cognition is our ability to understand and respond in a meaningful way to objects that we see, language that we hear, events that we experience, and any other stimuli in our environment. The ventral anterior temporal lobes (vATLs) play a critical role in such abilities^1–3^, but it remains unclear *how* neural activity in these regions encodes semantic information. Electrocorticography (ECoG) grid electrode data are a rare and informative resource, sensitive to activity at a wide range of frequencies including very high-frequency activity thought to indicate local neuronal activity^4–7^. Although prior work has shown that surface voltages express information about stimulus animacy (is it an animal or not? ^8,9^), voltages are a conflation of high- and lower-frequency activity; therefore, apparent representation of semantic information by the vATL may actually reflect incoming signals from more remote areas. We address these questions by applying regularised logistic regression — a multivariate decoding procedure that makes minimal assumptions about the nature of neural representations^10,11^ — to time-frequency power and phase extracted from a large sample of ECoG data (n = 18).

Extensive prior work suggests that the vATLs serve as a representational “hub”, interacting with multiple “spokes” that each encode modality-specific information about the environment. By virtue of these interactions, and over the course of learning and development, the vATLs come to express transmodal conceptual similarity structure and enable generalisation of knowledge across items and contexts^1–3,12^. The “hub-and-spoke” model is supported by convergent evidence from neuropsychology ^13^, positron emission tomography (PET^14^, distortion-corrected functional magnetic resonance imaging (fMRI^15–17^, magnetoencephalography (MEG^18^, neuro-computational simulations^3,9,19^, transcranial magnetic stimulation (TMS^20–22^, invasive cortical stimulation mapping^23,24^, and ECoG^25–28^.

Multivariate decoding approaches have only recently been applied to ECoG data in an attempt to characterise, not simply where and when task-related changes in neuronal activity occur, but what information those changes represent. Most of these analyses have focused on voltage^8,9,29–31^ (cf. ^32–34)^. However, voltage reflects a blend of activity at different frequencies and, while gamma (> 30 Hz) and high gamma (> ∼60 Hz) activity is thought to reflect local neuronal activity^4–7^, activity at lower frequencies may reflect transmission of information over varying cortical distances^35,36^, which may be important for assembling the spatially-distributed representations posited by the hub-and-spoke model.

The first aim of this paper was to clarify the role of time-frequency power and phase in semantic representation in the vATL and to relate this to voltage findings^8,9,29–31^ and to other sources of evidence^1^. The dataset consisted of ECoG data collected from a large cohort of patients (n = 18) while they named line drawings of animate and inanimate objects. ECoG offers excellent spatial and temporal resolution and the use of grid electrodes on the cortical surface offers much greater coverage than can be obtained with cortical depth electrodes — important for detecting semantic representations that may be distributed across space^6,9^. Crucially for this study, intracranial methods are necessary for revealing high gamma activity that is blocked by the skull and scalp when using noninvasive methods such as EEG or MEG^36,37^.

We used regularised logistic regression to decode information from time-frequency power and phase – a method that, unlike decoding methods that have previously been applied to ECoG time-frequency data^32–34^ makes very few assumptions about the properties of the underlying code^11^. Regularised regression can detect representations whether individual populations or frequencies encode pieces of information separately, or whether the relationship between information and neural activity can be understood only by considering multiple populations and/or frequencies simultaneously^11^. Since it relies on parameter fitting rather than simple correlations, it is resistant to false positives and false negatives associated with purely correlational approaches^29^.

We tested whether animacy (a) can be decoded from time-frequency power and/or phase; (b) if so, from which frequency ranges and at what timepoints; and (c) how these results relate to decoding of voltage. These analyses therefore allowed us to determine whether time-frequency- and voltage-based decoding offer similar or different conclusions about the timecourse of semantic representation in the vATL. Decoding accuracy in gamma and high gamma ranges (independently of other ranges) allowed us to determine whether semantic information is encoded by local neural activity within the vATL.

The second aim of this paper was to assess whether the dynamic and nonlinear changes to semantic representation observed in voltage decoding^9^ also arise within or across frequency ranges. Dynamic change in representation can be measured via the *temporal generalisation* of a decoding model^9,38–41^. The model is fitted to neural data at a given timepoint, then is assessed at other timepoints. If a classifier shows above-chance hold-out accuracy when fitted to timepoint A, and also shows above-chance accuracy when used to decode at timepoint B, we can conclude that information is encoded similarly at both timepoints. Conversely, if a classifier trained and succeeding in timepoint A then fails when tested on timepoint B, and if the reverse pattern also holds (a model trained on B succeeds on B but fails on A), this suggests that the target information is encoded by different neural patterns at different timepoints. That is, such a pattern suggests dynamic change in neural representation of the target information.

When applied to ECoG voltage data, this technique revealed four interesting characteristics of the semantic code in the vATL^9^:

1. Classifiers fitted and tested at overlapping 50 ms windows showed reliable decoding from about 100ms post stimulus onset through the remainder of the processing window (1650ms), suggesting that animacy is *constantly decodable* from the moment stimulus-driven activity reaches the vATL until the response.
2. Classifiers trained at a given window generalised to temporal neighbours (both preceding and succeeding), but not to temporally remote windows; that is, voltage-based neural decoding showed *local temporal generalisation*, suggesting a dynamically-changing neural code for animacy.
3. The width of the temporal generalisation window for a given classifier was quite narrow early in stimulus processing but widened over time, indicating that dynamic change to the voltage-based code is initially quite rapid but slows with further processing.
4. The direction of the neural code — whether an increase in voltage at a particular electrode signified increased or decreased probability that the stimulus is animate — changed over the course of processing, so that voltage deflections might signify different things at different timepoints at the same electrode.

Together these properties suggest that, while information about animacy is expressed in voltage throughout stimulus processing, the precise way that this information is encoded changes rapidly and nonlinearly, with change slowing as the system settles into a steady state. Moreover, these patterns mirrored those observed in a recurrent deep neural network model of semantic processing, suggesting that such models may offer a useful way of thinking about representational dynamics in real neural systems^9^.

The second goal of this paper was to assess whether these properties (constant decodability, local temporal generalisation, widening generalisation window, and change in code direction) are observed when decoding time-frequency power or phase information, across ranges or within each range considered independently. Answers to these questions could clarify whether these dynamic characteristics of representation arise from rapid local activity (i.e., driven by gamma/high gamma) and/or from more remote input (driven by lower frequencies).

## 1. Results

### 1.1. Relative decoding accuracy using spectral frequency power or phase vs. voltage

The ECoG data were collected from 6 - 52 subdural grid electrodes implanted over the ventral anterior temporal lobe (vATL) in 18 patients. The patients named line drawings of animals and inanimate objects that were used in prior work and matched for age of acquisition, concreteness, word frequency, and familiarity^42^. Time-frequency power and phase were extracted using complex Morlet wavelet convolution (see Methods). Classifiers were fitted using LASSO regularisation^43^ and assessed using ten-fold cross-validation. The classifiers were trained either on frequency features (vectors of power or phase values extracted from multiple frequencies from multiple electrodes at a single timepoint) or on voltage features (vectors of voltage values extracted from multiple electrodes from a 50 ms window centred on the timepoint of interest). Separate classifiers were trained for timepoints between 0 and 1650 ms in 10 ms time steps (0 ms, 10 ms, 20 ms, …). Each classifier was tested on every possible timepoint.

We first tested whether it was possible to decode the animacy of the stimuli using time-frequency power and/or phase and whether decoding performance was comparable to voltage^9^. We compared classifiers trained on frequency features of power or phase at 60 frequencies, logarithmically spaced between 4 and 200 Hz, to classifiers trained on voltage features. Figure 1A shows the results. Time-frequency power showed an almost identical decoding profile to voltage – hold-out accuracy rose to around 0.7 at 200 ms post stimulus onset and remained significantly above chance (0.5) throughout the time window. At no point was it possible to decode animacy from classifiers trained on phase.

**Figure 1:**
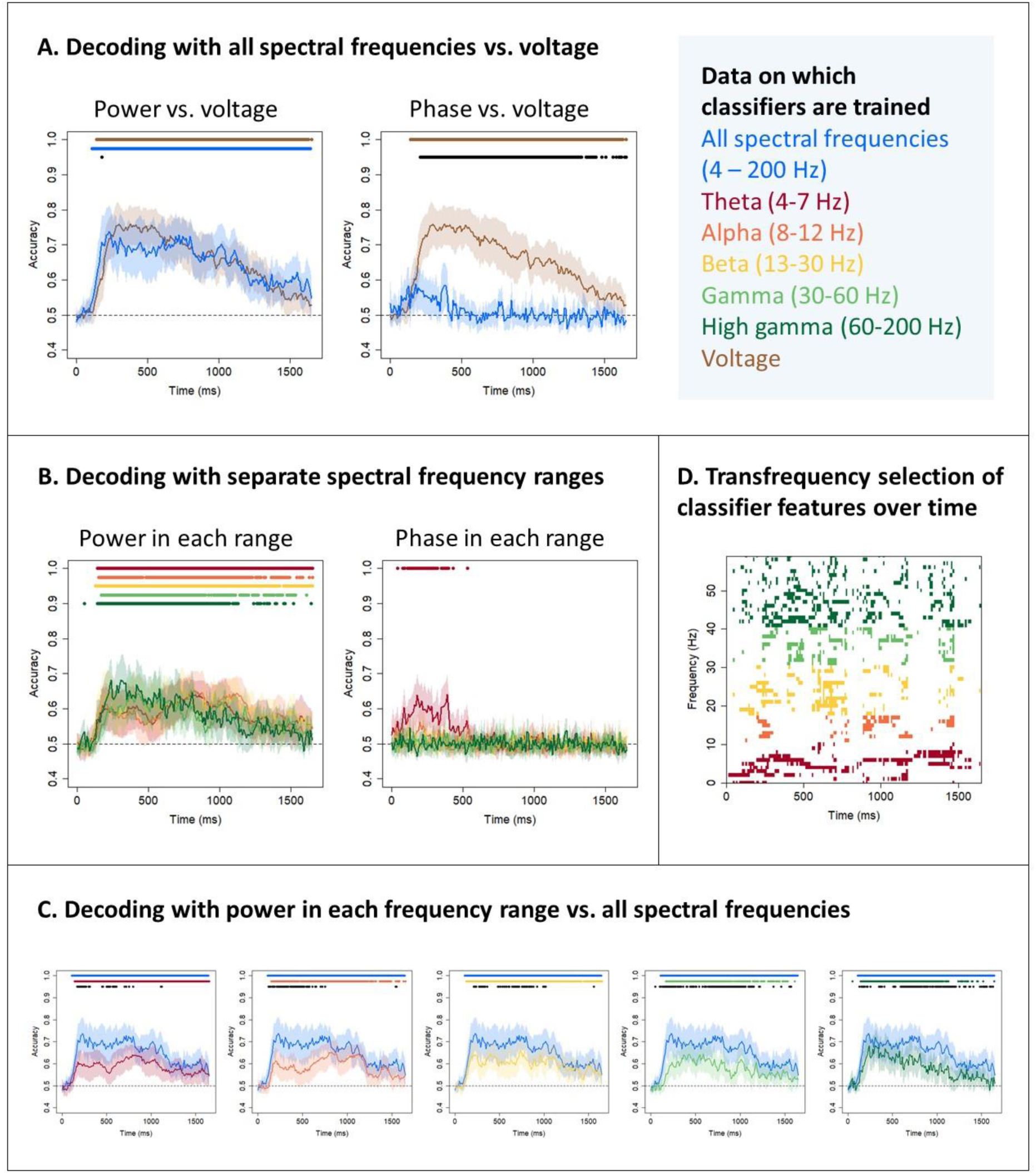
Semantic decoding with time-frequency power and phase. (A) Mean and 95% confidence interval of the hold-out accuracy for classifiers trained on power or phase frequency features for all frequencies between 4 and 200 Hz (blue) or on voltage features (brown). Coloured dots indicate a significant difference between classifier accuracy and chance (0.5, one-sample t-test with probabilities adjusted to control the false-discovery rate at α = 0.05). Black dots indicate a significant difference between accuracies at a given timepoint (paired t-tests with probabilities adjusted to control the false-discovery rate at α = 0.05). (B) Mean and 95% confidence interval of the hold-out accuracy for classifiers trained on frequency features composed only of power or phase values from a given range - theta (4 – 7 Hz, red), alpha (8 – 12 Hz, orange), beta (13 – 30 Hz, yellow), gamma (30 – 60 Hz, light green) and high gamma (60 – 200 Hz, dark green). Coloured dots indicate a significant difference between classifier accuracy and chance (0.5, one-sample t-test with probabilities adjusted to control the false-discovery rate at α = 0.05). (C) Mean and 95% confidence interval of the hold-out accuracy for classifiers trained on frequency features composed only of power or phase values from a given range compared to classifiers trained on frequency features including all frequencies between 4 and 200 Hz. Coloured dots indicate a significant difference between classifier accuracy and chance; black dots indicate a significant difference between accuracies. (D) Feature selection of each frequency at each timepoint (averaged across electrodes) for classifiers trained on frequency feature vectors including all frequencies between 4 and 200 Hz for a single patient. Features with a nonzero coefficient are shown in colour.

Next, to disentangle the role of local neural activity versus distal input, we trained classifiers on frequency features composed only of power or phase values from one frequency range - theta (4 – 7 Hz), alpha (8 – 12 Hz), beta (13 – 30 Hz), gamma (30 – 60 Hz) and high gamma (60 – 200 Hz). Figure 1B shows the decoding profile in each range. For power, the patterns were almost indistinguishable, showing the same rise to about 0.6 and remaining above chance for most of the time window of interest. On the assumption that gamma and high gamma frequencies represent local neural activity, these results confirmed that local neural activity in the vATL expresses semantic information. However, we found that power in every range was sufficient, and no range was necessary, for above-chance decoding. For phase, decoding was rarely above chance, with the exception of weak decoding using theta data in the window 0 - 500 ms. The remaining analyses therefore focused on characterising semantic information encoded in time-frequency power.

Figure 1C shows hold-out accuracy for classifiers trained on power within a single range compared to accuracy for classifiers trained on power at all 60 frequencies. For each range there was a time window during which decoding on all frequencies reached its peak and decoding on a single range performed significantly less well. This indicated that no individual range contains sufficient information to enable decoding accuracy comparable to accuracy based on all frequencies. To investigate which combination of frequencies enabled such successful decoding, we inspected the coefficients on each frequency. Figure 1D shows these averaged across electrodes for one example patient (similar plots for all patients are shown in Supplementary Materials S1, Figure S1; plots showing coefficients on individual electrodes for the one sample patient are shown in Supplementary Materials S1, Figure S2). Throughout the time window, frequencies across the whole range were selected by the classifier and thus contributed to decoding performance. This is evidence that the vATL represents semantic information via a *transfrequency* code, by which we mean that frequencies within the theta, alpha, beta, gamma and high gamma ranges all contribute to information representation.

### 1.2. Does the time-frequency semantic code change dynamically?

Our second question was whether the time-frequency code changes dynamically over stimulus processing in a manner similar to the voltage code. The preceding results show that time-frequency power exhibits one property also observed in voltage, namely constant decodability - animacy can be decoded from whole-spectrum power, and from power within each frequency range independently, from about 100-200ms post onset until about the mean onset of naming (1190ms in the subset of patients analysed by Rogers et al.^9^). Next, we applied the method of temporal generalisation to investigate whether time-frequency power, like voltage, shows (a) local temporal generalisation, indicating a changing neural code, and (b) a widening generalisation window, indicating rapid initial change that slows over time^9^.

Figure 2A shows the generalisation profile of classifiers trained on power at all frequencies (left) and those trained on voltage (right). The diagonal of each matrix indicates the hold-out accuracy of classifiers trained and tested at the same timepoint. Points closer to the diagonal indicate how well a classifier generalises to neighbouring timepoints, while points far from the diagonal indicate how well it generalises to timepoints further away. Local temporal generalisation is signified by better decoding close to the diagonal. In these plots, the most saturated colours indicate classifier accuracies statistically indistinguishable from the best-performing classifier at the same timepoint. Increasingly desaturated colours indicate accuracies that are reliably worse than the best-performing classifier at increasingly strict significance thresholds. Horizontal black bars indicate classifiers that did not exceed chance performance even when evaluated on held-out stimuli from the training timepoint.

**Figure 2:**
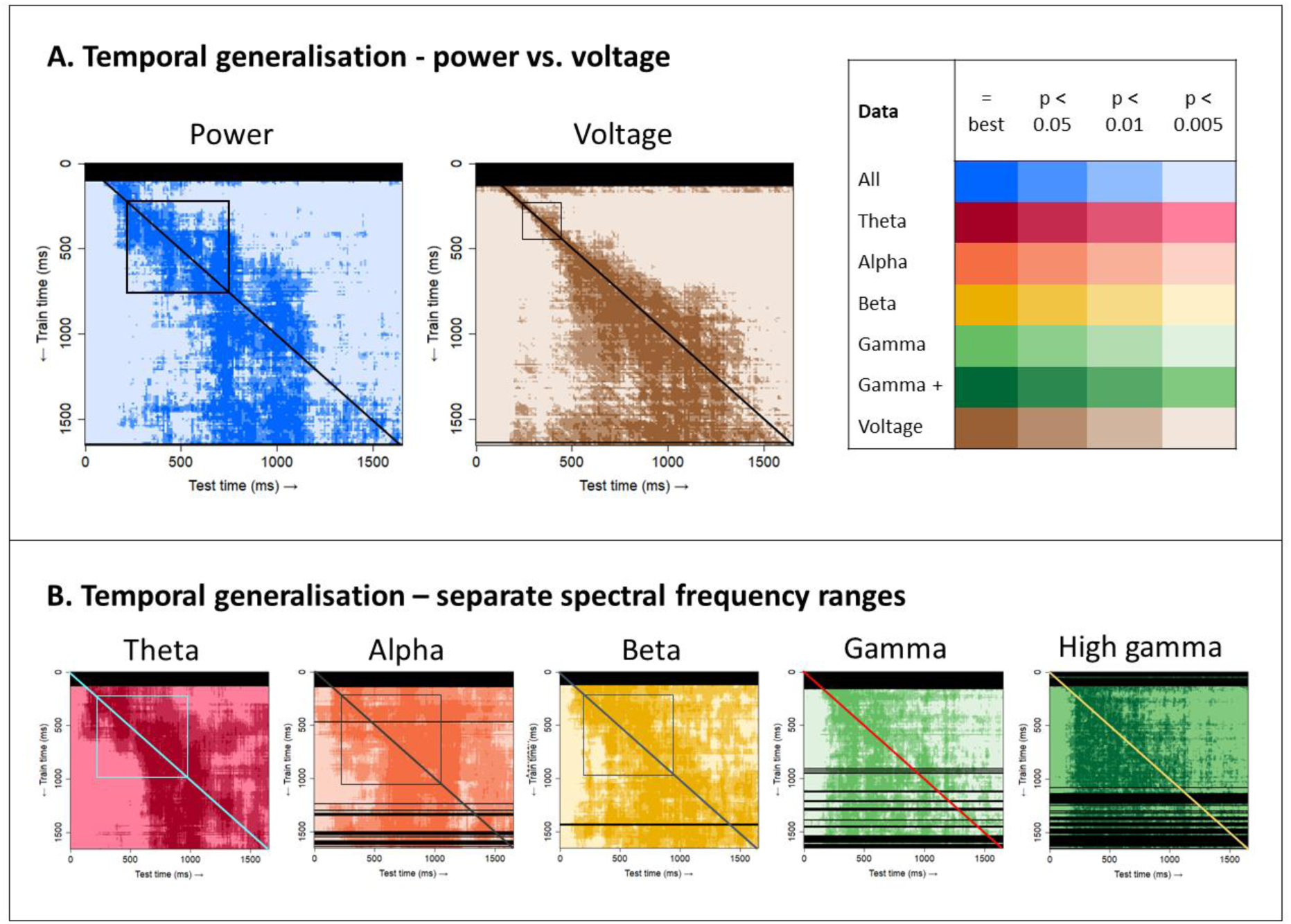
Temporal generalisation of classifiers. For each test time, the train time (vertical dimension) of the best classifier, plus the train times of other classifiers that perform statistically indistinguishably from the best (p > 0.05, paired t-test with probabilities adjusted to control the false-discovery rate at α = 0.05), are shown in the most saturated colour. Progressively desaturated colours show the train time of classifiers exhibiting a significant difference from the best classifier (p < 0.05, p < 0.01, p < 0.005). Timepoints where classifiers trained at that timepoint do not perform significantly better than chance even when tested at the training timepoint (0.5, one-sample t-test with probabilities adjusted to control the false-discovery rate at α = 0.05) are shown in black. Local temporal generalisation is indicated by significantly better performance on timepoints at or near the train time (the diagonal shown on each plot) than at more remote timepoints. The squares highlight cases where hold-out accuracies at two training timepoints (the two corners on the diagonal) are both significantly better than when the same classifiers are used to decode more remote timepoints (the two off-diagonal corners). (A) Results for classifiers trained on power frequency features for all frequencies between 4 and 200 Hz (blue) or on voltage features (brown). (B) Results for classifiers trained on power frequency features from a single range - theta (4 – 7 Hz, red), alpha (8 – 12 Hz, orange), beta (13 – 30 Hz, yellow), gamma (30 – 60 Hz, light green) and high gamma (60 – 200 Hz, dark green).

The superimposed squares on each plot provide illustrative referents for evaluating whether the temporal generalisation pattern signifies a dynamically changing code. If classification accuracy is significantly higher for the two on-diagonal corners than the two off-diagonal corners, this signifies that (a) the neural signal reliably encodes semantic information at the two timepoints but (b) the signal changes in nature – whatever features of the signal support the good performance at the training timepoint are not present at the more remote testing points. This pattern is clearly observed for both voltage and whole-spectrum power (Figure 2A), indicating that the transfrequency semantic code exhibits dynamic properties similar to voltage.

When decoding from each range separately, however, the local generalisation pattern was less clear overall, and arguably present only in lower frequency ranges (theta, alpha, and beta; Figure 2B). Gamma and high gamma showed a different pattern: classifiers fitted at a given timepoint generalised approximately equally well to all prior timepoints back to the first timepoint where decoding was reliable (about 100-200ms post stimulus onset). That is, classifiers trained on gamma and high gamma showed excellent *retrograde* generalisation. This pattern yields the vertical “edge” clearly evident in the high gamma matrix, and blurrier but still present in the gamma matrix. The classifiers also generalised to future timepoints, but with performance often tailing off for more remote future timepoints — that is, they showed diminishing *anterograde* generalisation.

Together these observations suggest two interesting things about the semantic code as expressed within and across different frequency ranges. First, whatever information a classifier trained on gamma or high gamma power exploits at its own training timepoint must also be present in the signal at all earlier time points, all the way back to the first moment semantic information appears (producing good retrograde generalisation). However, information exploited by a classifier at its training timepoint may *not* be present in the signal at future timepoints, suggesting that these features “drop out” over the course of stimulus processing, producing a graded drop-off in anterograde generalisation. Second, the dynamically-changing pattern observed in the transfrequency code may be carried predominantly by lower frequency information.

We decided to focus subsequent analyses on frequency ranges in which a dynamic code was most clearly observed, namely voltage and whole-spectrum power. We next assessed whether the window of above-chance temporal generalisation widened over time, consistent with an overall slowing of representational change. To this end, we grouped the classifiers into ten clusters based on similarity in their temporal decoding profiles (see Methods). That is, classifiers that show similar patterns of prediction accuracy across all test timepoints were clustered together, and we computed the mean accuracy profile over time for each cluster. Figure 3 shows the result. Classifiers trained earlier showed a sharper rise and fall in accuracy, whereas classifiers trained later exhibited a wider window of generalisation (similar plots for classifiers trained on separate frequency ranges are shown in Supplementary Materials 1, Figure S3; statistics supporting this claim are reported in Supplementary Materials 1, Figure S4). Thus both voltage and trasnfrequency power show a slowing of dynamic representational change over the course of stimulus processing.

**Figure 3:**
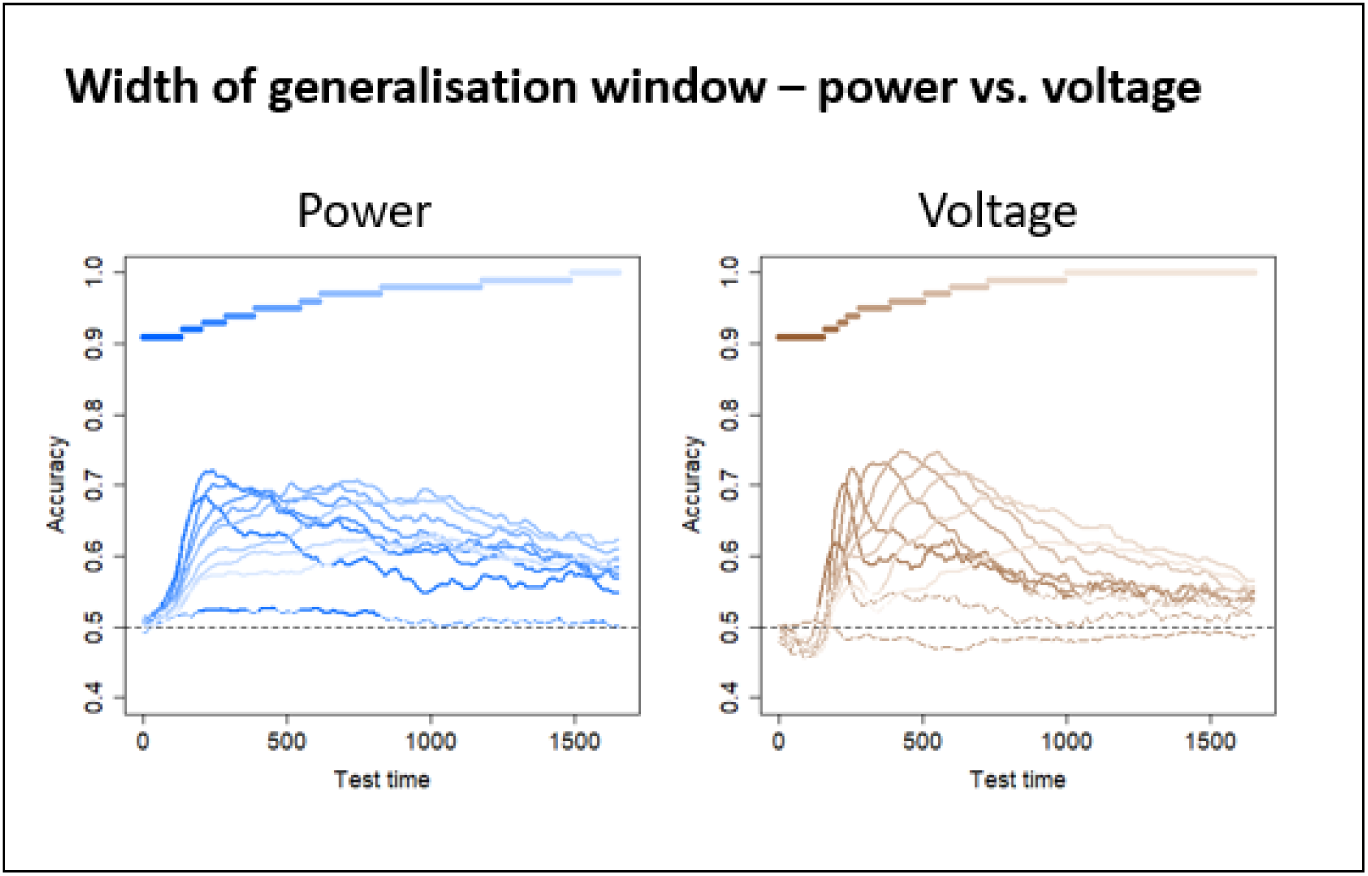
Width of generalisation window. Classifiers trained on power frequency features for all frequencies between 4 and 200 Hz (shades of blue) or on voltage features (shades of brown) were grouped into ten clusters via agglomerative hierarchical clustering (see Methods). The “timecourses” show the mean hold-out accuracy for classifiers within each cluster at each timepoint. Lines are solid where there was a significant difference between classifier accuracy and chance (0.5, one-sample t-test with probabilities adjusted to control the false-discovery rate at α = 0.05) and dashed where there is no significant difference. Coloured bars show the grouped timepoints in each cluster. Statistics supporting this pattern are reported in Supplementary Materials 1, Figure S4).

Finally, we assessed whether the observed dynamic change was accompanied by change in the apparent code direction—that is, whether an increase in voltage or in power predicted a greater likelihood that the stimulus is an animal at some points in time, and a greater likelihood that it is inanimate at others. Change in code direction is another hallmark of nonlinear representational change^9^. To see this, we aggregated the classifier coefficients across patients for each timepoint and plotted these on the cortical surface. Specifically, at each cortical location, we first calculated the number of electrodes across patients that had a non-zero coefficient, then computed the proportion of these having a negative coefficient (since animals were coded as 0 and inanimate objects as 1, a negative coefficient means that an increase in voltage or power is associated with increased probability that the stimulus is an animal). Figure 4 shows these proportions at 4 different timepoints post stimulus onset, for classifiers trained on whole-spectrum power (top) or voltage (bottom). Warm colours indicate areas and timepoints where most selected electrodes have negative coefficients, while cool colours indicate areas and timepoints where most selected electrodes have positive coefficients. Green shades indicate areas and timepoints where selected electrodes have a mixture of positive and negative coefficients. We also animated the coefficient change over time, with videos for classifiers trained on all frequencies, individual frequency ranges, and voltage available at https://github.com/slfrisby/ECoG_LASSO/.

**Figure 4:**
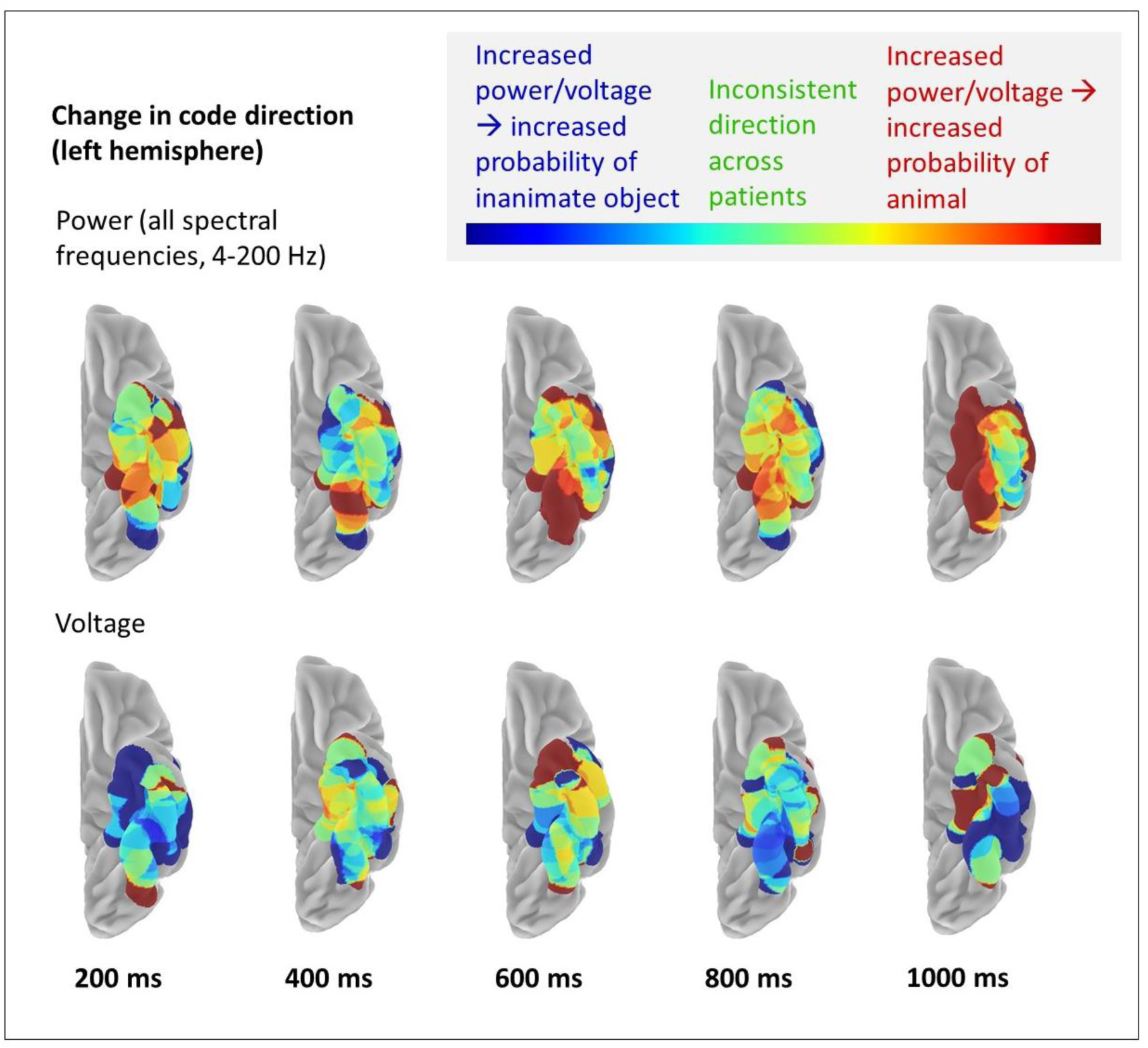
Change in code direction. Coefficients for classifiers trained on power frequency features for all frequencies between 4 and 200 Hz, or on voltage features, are projected onto the ventral surface of the left hemisphere. Time is shown relative to stimulus onset. Colours indicate the proportion of classifier coefficients at that location with the same sign (see Methods). Warm colours indicate that coefficients are mostly negative across patients (since animals were coded as 0 and inanimate objects as 1, a negative coefficient indicates that an increase in power is associated with increased probability that the stimulus is animate); cool colours indicate that coefficients are mostly positive across patients (an increase in power is associated with increased probability that the stimulus is inanimate); green shades indicate that the region was selected in multiple patients but that the sign of the coefficient is not consistent across patients.

For both voltage and power, changes in code direction were observed. For voltage, for instance, the anterior-most part of the ventral temporal lobe was uniformly positive at 200ms and uniformly negative at 600ms. By 1000ms, midway between the anterior and posterior extent of coverage, coefficients showed a medial-to-lateral change in direction, suggesting that voltage increases signify animals in the more medial portion and non-animals in the more lateral portion. Whole-spectrum power also showed changes in code direction over time; for instance, in Figure 4, the medial anterior portion of ventral temporal cortex showed a negative relationship between power and animacy at 400ms, but a positive relationship by 1000ms. At 1000ms, the transfrequency coefficients also showed a medial-to-lateral change, with increased power signifying animals all along the medial extent of the coverage area, but showing a more mixed pattern across the more lateral extent.

## 2. Discussion

While it has long been clear that the vATLs play an important role in human semantic cognition^1–3^, precisely *how* neural populations encode such information remains uncertain. Recent studies applying multivariate decoding to voltages recorded from the cortical surface suggest that semantic information is expressed in a rapidly-changing code^9^, but, with respect to uncovering local neural representations, such studies remain limited, since voltages can be influenced by a variety of sources both local and remote. The current work instead applied multivariate decoding to a time-frequency decomposition of ECoG data, with two results that advance our understanding of the vATL semantic code in important ways. First, information about animacy is expressed by power in all frequencies simultaneously - it appears within each range considered independently, and jointly across all ranges, for most of the duration of stimulus processing. Moreover, decoding accuracy was reliably better when models were fitted to all frequencies simultaneously compared to each individual range; and classifiers fitted to the whole spectrum always selected features from all frequency ranges no matter which timepoint they were trained on. This result strongly suggests conjoint encoding of semantic information across the power spectrum^11^. Second, decoding from both power and voltage revealed characteristics of dynamic representational change: constant decodability, local temporal generalisation, a widening generalisation window, and changes in code direction. This is likely to be driven by low frequencies – gamma and high gamma classifiers showed a contrasting pattern of excellent retrograde generalisation back to the earliest successful decoding timepoint, but anterograde generalisation that diminished for more remote timepoints.

Assuming that the frequency of activity is inversely related to the distance over which information is being transmitted^35,36^, our results are consistent with the claim that both local neuronal activity within the vATL (reflected in gamma and/or high gamma power) and distal inputs to the region (reflected in theta, alpha and beta) encode semantic information. These findings therefore agree with multiple univariate studies that variously associate theta^44^, alpha^45^, beta^27,46^, gamma and high gamma^34,47–58^ with semantic tasks. One interpretation of our results is that information is represented redundantly in multiple frequencies. A second possibility is that, since the brain exhibits properties of a dynamical system^59,60^, rhythmic patterns may be an epiphenomenon of normal functioning or even of epileptiform activity^61–63^. Therefore, some frequency ranges may represent information not in a “strong” sense (meaning that the brain uses time-frequency power to represent information for downstream computation) but in a “weak” sense (meaning that time-frequency power is decodable but is not used as the principal means of representation^64^).

However, the finding that classifiers trained on whole-spectrum power outperform those trained on a single frequency range and select features from across the frequency spectrum (Figures 1C and 1D) supports a more nuanced conclusion. Classifiers trained with LASSO regularisation prefer sparse solutions, in which only a small subset of features receive nonzero coefficients. If the information in the different frequency ranges were redundant, a classifier would not assign nonzero coefficients to many features that spanned the frequency range. Instead, these results support the notion of *transfrequency* representation – information is encoded *across* multiple frequency ranges (sometimes called frequency “multiplexing”^64^). Some previous univariate studies have proposed specific roles that different frequency ranges may play - abstraction from multiple episodic experiences^65^, binding multiple features of an item^6,65–68^, encoding semantic similarity^69,70^, and enabling retrieval independent of sensory modality^65^ in a way that is appropriate for the context^71–74^. However, on the simple assumption that the frequency of activity is inversely related to transmission distance, the notion of transfrequency representation is highly consistent with the hub-and-spoke model^1–3^. According to this model, neither the vATL “hub” nor the modality-specific “spokes” are individually sufficient for full semantic cognition – rather, information is encoded conjointly in multiple regions and in the interactions between them^11^.

This study was the first to investigate whether activity in different frequency ranges exhibits the same pattern of dynamic representational change seen in voltage^9^. Although dynamic change was clearly evident when decoding whole-spectrum power, gamma and high gamma showed a different pattern – excellent retrograde generalisation and diminishing anterograde generalisation. The good retrograde generalisation indicates that features exploited by a classifier at its training timepoint must also be present, and must signify the same information in the same way, for all preceding timepoints (up to the earliest successful timepoint). That is, this part of the code must be relatively *static* over processing from the earliest arrival of perceptual signals up to the time where the classifier is fitted. Conversely, the diminishing anterograde generalisation indicates that features exploited by a classifier at its training timepoint eventually fall away at future timepoints.

Figure 5 illustrates in schematic form what such a decoding pattern might suggest about how animacy is encoded in high-frequency activity (and thus, by inference, in local neural activity). Each curve represents a hypothetical response to animate stimuli in the gamma or high gamma range for one electrode over time. Every electrode shows increased gamma power for animate stimuli from early in processing, but the timecourse of this change differs between electrodes. Where gamma increases most strongly early in processing, it also diminishes most rapidly; where gamma increases are smaller initially, they persist longer. Since the LASSO penalty prefers to assign features the smallest coefficients possible, models fitted with LASSO regularisation will select whichever electrodes show the strongest response at the training tinepoint. Thus models fitted to the earliest timepoints will select the red electrodes, those fitted slightly later will select the orange ones, etc. Generalisation to future timepoints will diminish as the gamma power decreases on the selected electrodes, producing diminishing anterograde generalisation. Because all electrodes carry useful signal from the onset, however, each classifier generalises successfully to all prior time points, producing good retrograde generalisation. The generalisation pattern is thus consistent with the view that animacy is encoded in the local activity of many neural populations, some responding strongly but then quickly diminishing, and other initially responding less strongly but persisting for a longer duration.

**Figure 5:**
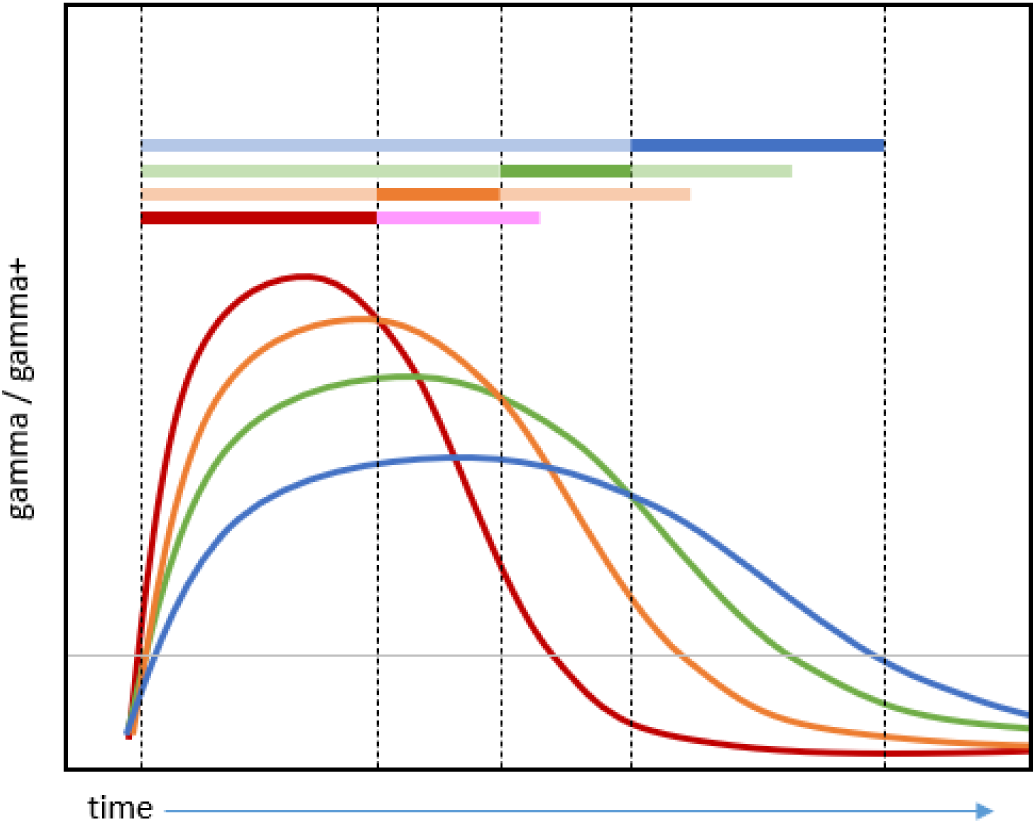
Hypothesised semantic code in gamma and/or high gamma producing full retrograde generalisation and limited anterograde generalisation. Each curve represents the hypothesised gamma/high gamma response to animate stimuli by one electrode. Where curves are above the grey horizontal line, they encode information about animacy. All electrodes encode animacy from early in processing, but models regularised with the LASSO penalty will place coefficients on whichever electrode has the strongest response at a given timepoint. The saturated parts of the coloured horizontal bars thus indicate which electrode is selected by a classifier at each timepoint. The desaturated parts indicate the generalisation profile of the resulting classifier. Electrodes that show the strongest responses early on diminish in power most rapidly, limiting anterograde generalisation. Because all electrodes carry information about animacy from the onset of the signal, however, all classifiers generalise well to the beginning of the processing window.

Finally, the current work carries important implications for interpreting other sources of information about semantic representation. Noninvasive methods such as EEG and MEG cannot discern activity within high gamma frequencies because it is blocked by the skull and the scalp. The current work suggests that dynamic representational change is mainly observed in lower frequency ranges that are more readily measurable via these other techniques—thus the comparatively static code revealed in high frequency ranges may be obscured in other imaging modalities. Additionally, the current work highlights circumstances in which representations in neural activity differ in their nature from representations observed in neuro-computational models (which exhibit the dynamic properties observed in voltage and transfrequency power, not the comparatively static code observed in high frequencies). Further work should explore the circumstances under which representational properties in brains and machines converge and diverge^75^.

### 2.1. Conclusion

To summarise, we have shown that it is possible to decode semantic information from every frequency range between theta and high gamma. Although individual frequency ranges contained enough information to drive decoding above chance, decoding from the whole frequency spectrum at once produced the highest decoding accuracy, equivalent to using voltage. Decoding from the whole frequency spectrum also revealed the same set of dynamic properties observed in voltage; however decoding of high-frequency ranges alone showed a qualitatively different pattern, suggesting that local firing activity expresses a more static semantic code. Together these results provide an important step toward understanding how neural systems encode semantic information.

## 3. Online methods

### 3.1. Patients

19 patients participated in the study (labelled 01-22 to enable comparison with previous work^8,9,28,29^). Of these 19 patients, one was later excluded because too many trials were contaminated with artefacts (see Preprocessing – ECoG), leaving 18 patients for decoding. For one further patient (Patient 20), no exact MNI coordinates were available for their electrodes. This patient was included in the analyses, but excluded from visualisation (see Experimental Questions). All patients were native speakers of Japanese. Information about patients’ age, sex, handedness, and clinical presentation is summarised in Table 1.

**Table 1:**
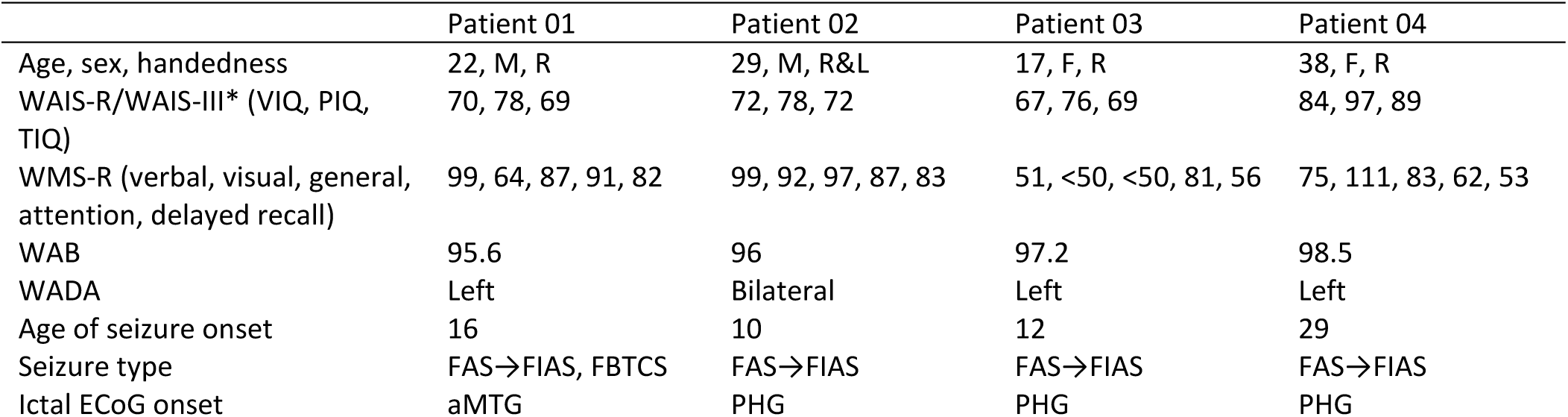
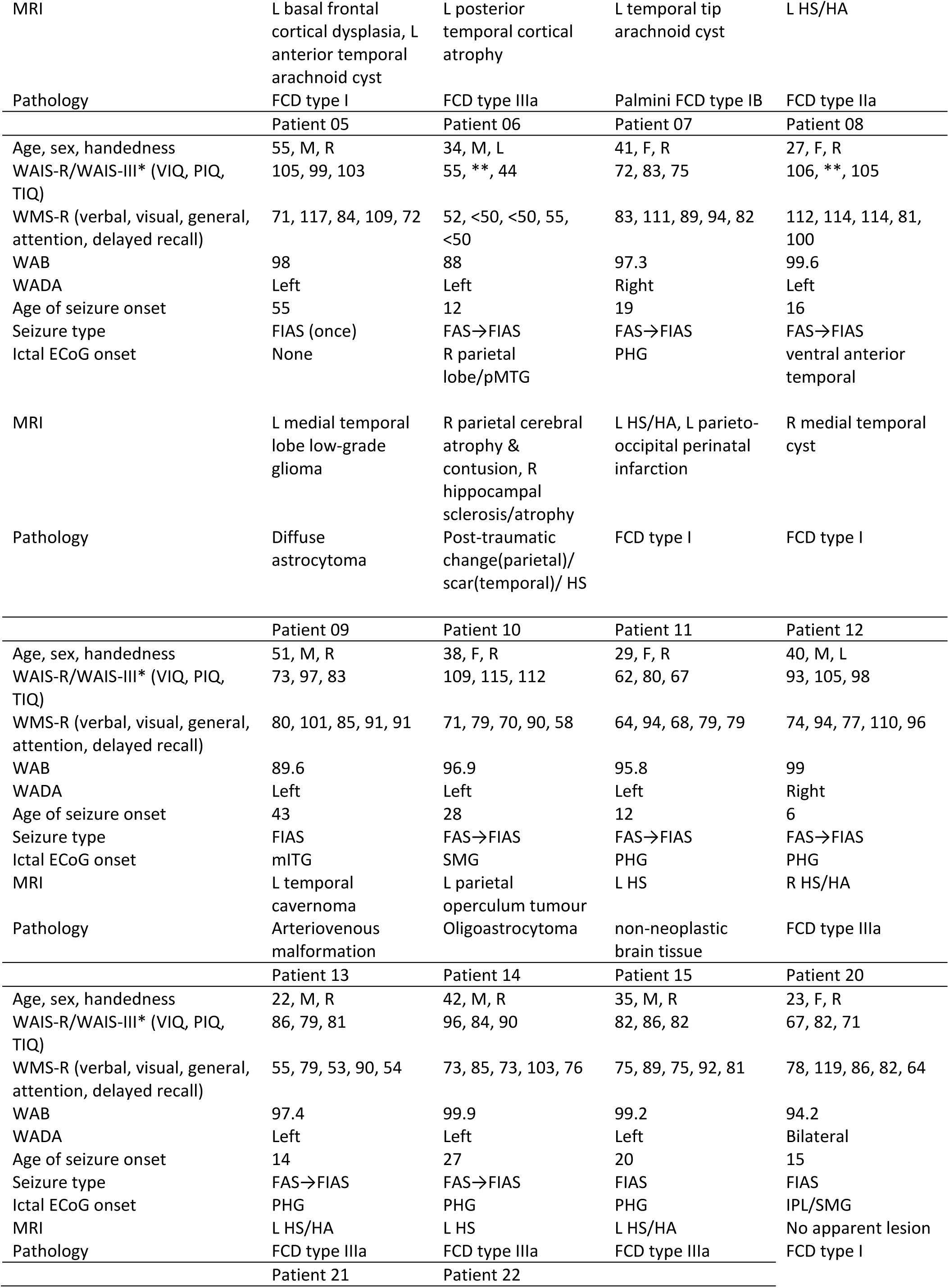
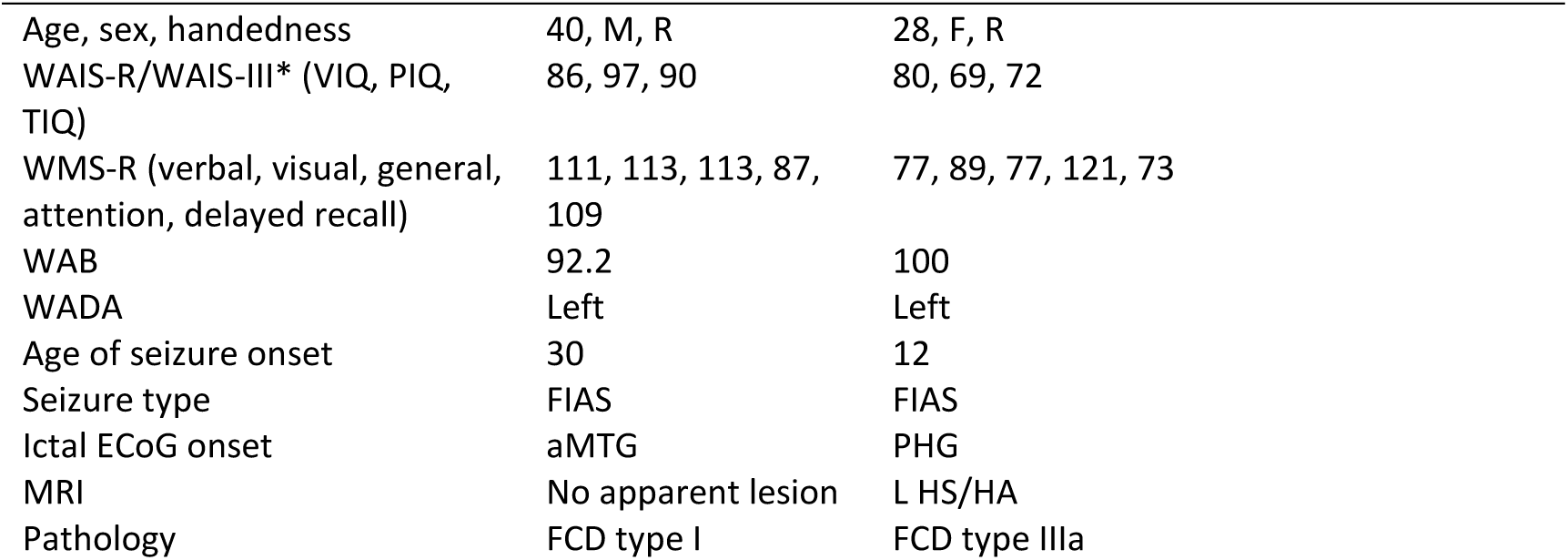
Patient characteristics. Abbreviations: WAIS-R – Wechsler Adult Intelligence Scale (1991), WAIS-III - Wechsler Adult Intelligence Scale (1997), VIQ – Verbal IQ, PIQ – Performance IQ, TIQ – full-scale IQ, WMS-R – Wechsler Memory Scale (1987), WAB – Western Aphasia Battery, FAS – focal aware seizure, FIAS – focal impaired awareness seizure, FBTCS – focal to bilateral tonic-clonic seizure, aMTG – anterior middle temporal gyrus, PHG – parahippocampal gyrus, pMTG – posterior middle temporal gyrus, mITG – medial inferior temporal gyrus, SMG - supramarginal gyrus, ITG – inferior temporal gyrus, IPL – intraparietal lobule, HS – hippocampal sclerosis, HA – hippocampal atrophy, FCD – focal cortical dysplasia. * - The WAIS-R was used to test patients 01-06 and the WAIS-III was used to test other patients. ** - missing score.

|  | Patient 01 | Patient 02 | Patient 03 | Patient 04 |
| --- | --- | --- | --- | --- |
| Age, sex, handedness | 22, M, R | 29, M, R&L | 17, F, R | 38, F, R |
| WAIS-R/WAIS-III* (VIQ, PIQ, TIQ) | 70, 78, 69 | 72, 78, 72 | 67, 76, 69 | 84, 97, 89 |
| WMS-R (verbal, visual, general, attention, delayed recall) | 99, 64, 87, 91, 82 | 99, 92, 97, 87, 83 | 51, <50, <50, 81, 56 | 75, 111, 83, 62, 53 |
| WAB | 95.6 | 96 | 97.2 | 98.5 |
| WADA | Left | Bilateral | Left | Left |
| Age of seizure onset | 16 | 10 | 12 | 29 |
| Seizure type | FAS→FIAS, FBTCs | FAS→FIAS | FAS→FIAS | FAS→FIAS |
| Ictal ECoG onset | aMTG | PHG | PHG | PHG |
| MRI | L basal frontal cortical dysplasia, L anterior temporal arachnoid cyst | L posterior temporal cortical atrophy | L temporal tip arachnoid cyst | L HS/HA |
| Pathology | FCD type I | FCD type IIIa | Palmini FCD type IB | FCD type IIa |
|  | Patient 05 | Patient 06 | Patient 07 | Patient 08 |
| Age, sex, handedness | 55, M, R | 34, M, L | 41, F, R | 27, F, R |
| WAIS-R/WAIS-III* (VIQ, PIQ, TIQ) | 105, 99, 103 | 55, **, 44 | 72, 83, 75 | 106, **, 105 |
| WMS-R (verbal, visual, general, attention, delayed recall) | 71, 117, 84, 109, 72 | 52, <50, <50, 55, <50 | 83, 111, 89, 94, 82 | 112, 114, 114, 81, 100 |
| WAB | 98 | 88 | 97.3 | 99.6 |
| WADA | Left | Left | Right | Left |
| Age of seizure onset | 55 | 12 | 19 | 16 |
| Seizure type | FIAS (once) | FAS→FIAS | FAS→FIAS | FAS→FIAS |
| Ictal ECoG onset | None | R parietal lobe/pMTG | PHG | ventral anterior temporal |
| MRI | L medial temporal lobe low-grade glioma | R parietal cerebral atrophy & contusion, R hippocampal sclerosis/atrophy | L HS/HA, L parieto-occipital perinatal infarction | R medial temporal cyst |
| Pathology | Diffuse astrocytoma | Post-traumatic change(parietal)/scar(temporal)/ HS | FCD type I | FCD type I |
|  | Patient 09 | Patient 10 | Patient 11 | Patient 12 |
| Age, sex, handedness | 51, M, R | 38, F, R | 29, F, R | 40, M, L |
| WAIS-R/WAIS-III* (VIQ, PIQ, TIQ) | 73, 97, 83 | 109, 115, 112 | 62, 80, 67 | 93, 105, 98 |
| WMS-R (verbal, visual, general, attention, delayed recall) | 80, 101, 85, 91, 91 | 71, 79, 70, 90, 58 | 64, 94, 68, 79, 79 | 74, 94, 77, 110, 96 |
| WAB | 89.6 | 96.9 | 95.8 | 99 |
| WADA | Left | Left | Left | Right |
| Age of seizure onset | 43 | 28 | 12 | 6 |
| Seizure type | FIAS | FAS→FIAS | FAS→FIAS | FAS→FIAS |
| Ictal ECoG onset | mITG | SMG | PHG | PHG |
| MRI | L temporal cavernoma | L parietal operculum tumour | L HS | R HS/HA |
| Pathology | Arteriovenous malformation | Oligoastrocytoma | non-neoplastic brain tissue | FCD type IIIa |
|  | Patient 13 | Patient 14 | Patient 15 | Patient 20 |
| Age, sex, handedness | 22, M, R | 42, M, R | 35, M, R | 23, F, R |
| WAIS-R/WAIS-III* (VIQ, PIQ, TIQ) | 86, 79, 81 | 96, 84, 90 | 82, 86, 82 | 67, 82, 71 |
| WMS-R (verbal, visual, general, attention, delayed recall) | 55, 79, 53, 90, 54 | 73, 85, 73, 103, 76 | 75, 89, 75, 92, 81 | 78, 119, 86, 82, 64 |
| WAB | 97.4 | 99.9 | 99.2 | 94.2 |
| WADA | Left | Left | Left | Bilateral |
| Age of seizure onset | 14 | 27 | 20 | 15 |
| Seizure type | FAS→FIAS | FAS→FIAS | FIAS | FIAS |
| Ictal ECoG onset | PHG | PHG | PHG | IPL/SMG |
| MRI | L HS/HA | L HS | L HS/HA | No apparent lesion |
| Pathology | FCD type IIIa | FCD type IIIa | FCD type IIIa | FCD type I |
|  | Patient 21 | Patient 22 |  |  |

Table 1: Patient characteristics.
|  |  |  |
| --- | --- | --- |
| Age, sex, handedness | 40, M, R | 28, F, R |
| WAIS-R/WAIS-III* (VIQ, PIQ, TIQ) | 86, 97, 90 | 80, 69, 72 |
| WMS-R (verbal, visual, general, attention, delayed recall) | 111, 113, 113, 87, 109 | 77, 89, 77, 121, 73 |
| WAB | 92.2 | 100 |
| WADA | Left | Left |
| Age of seizure onset | 30 | 12 |
| Seizure type | FIAS | FIAS |
| Ictal ECoG onset | aMTG | PHG |
| MRI | No apparent lesion | L HS/HA |
| Pathology | FCD type I | FCD type IIIa |

Each patient was implanted with subdural electrodes arranged in grids or strips for presurgical monitoring (mean 88 electrodes, range 56 – 108 electrodes per patient). 15 patients had electrodes implanted in the left hemisphere, of which 10 – 42 electrodes (mean 25 electrodes) were within grids or strips that covered the vATL (where grids or strips included both electrodes that covered ventral anterior temporal cortex and electrodes that covered lateral or posterior temporal cortex, the whole grid or strip was included in the analysis). In the remaining 3 patients, with electrodes implanted in the right hemisphere, 6 – 28 electrodes (mean 20 electrodes) were within grids or strips that covered the vATL. Grids in which no electrodes covered the vATL were excluded from analysis. Electrodes that were included in the analysis are shown in Figure 6. The electrodes were platinum, with a recording diameter of 2.3 mm and an inter-electrode distance of 1 cm (ADTECH, WI). Four patients were also implanted with depth electrodes (sEEG), but these were not included in the analysis.

**Figure 6:**
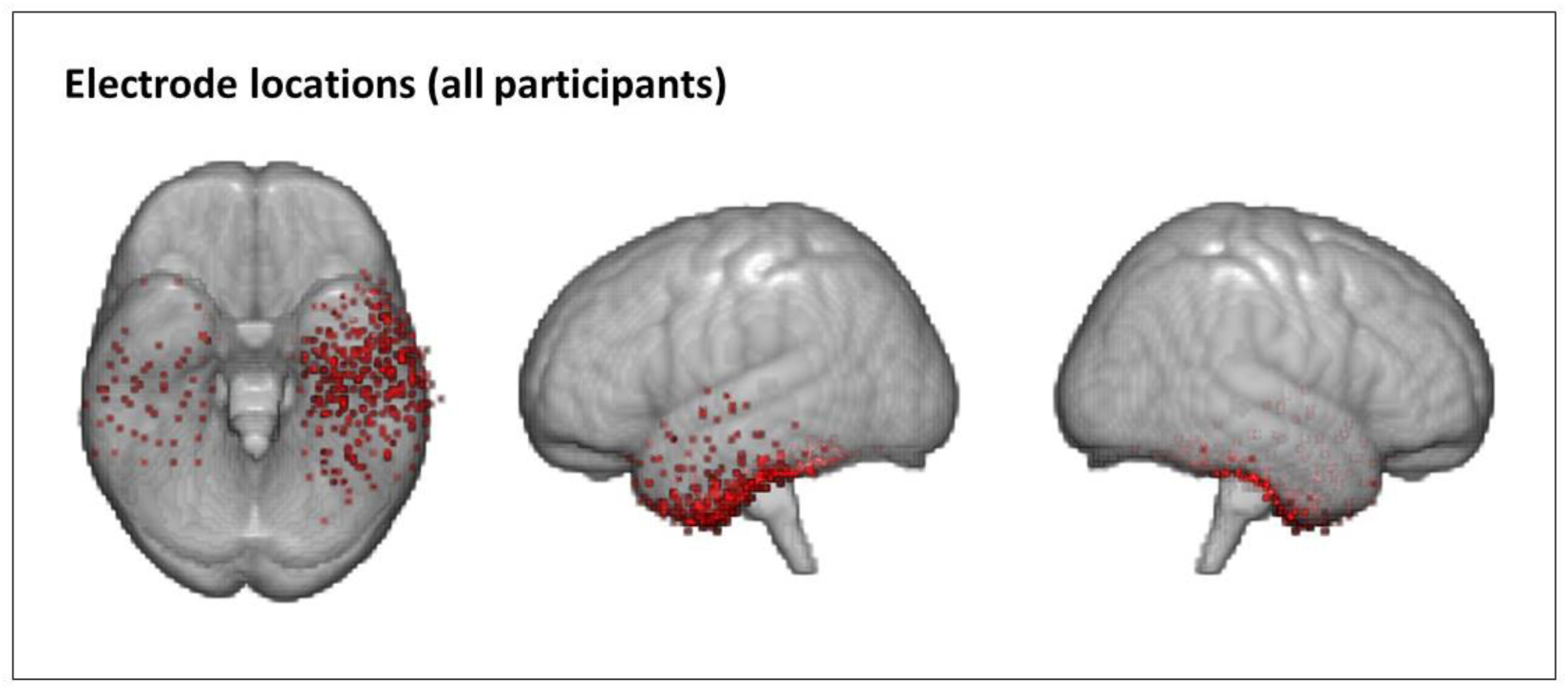
Electrode locations for all patients. Electrodes are overlaid on an MNI template (MNI152NLin2009cAsym).

All patients gave written informed consent and the study was approved by the ethics committee of the Kyoto University Graduate School of Medicine (#C533).

### 3.2. Stimuli, task, and acquisition

Stimuli were the same 100 line drawings used previously^8,9,28^ – 50 animals and 50 nonliving items including buildings, tools, musical instruments and other household objects^76^; https://github.com/slfrisby/ECoG_LASSO/tree/main/data_info/stimuli/). There were no significant differences between the categories with respect to age of acquisition, visual complexity, concreteness, familiarity, word frequency, name agreement and non-semantic visual structure^9,77^. MATLAB r2010a was used to display stimuli on a PC screen.

There were four runs per patient, collected in a single session. Within each run, each stimulus was presented once in a random order. Stimuli were presented for 5 seconds each, with no break between stimuli.

Patients were instructed to name each picture as quickly and accurately as possible. Data for nine patients were recorded at 2000 Hz (with a low-pass filter of 600 Hz) and data for ten patients were recorded at 1000 Hz (with a low-pass filter of 300 Hz). Time of naming onset was measured. Responses and eye fixation were monitored via video.

### 3.3. Data analysis

#### 3.3.1. Preprocessing – structural MRI

A clinical MPRAGE T1-weighted anatomical scan was acquired before and after electrode implantation. The location of each electrode was identified on each 2D slice of the post-surgical scan. *fnirt* in FSL (https://fsl.fmrib.ox.ac.uk/fsl/fslwiki/^78,79^) was used to coregister the electrode positions to the pre-surgical scan and then to MNI space (MNI152NLin2009cAsym). The position of each electrode was then manually adjusted to the surface.

#### 3.3.2. Preprocessing – ECoG

Preprocessing was performed using in-house MATLAB (r2023b) code, composed of functions from EEGLAB (2023.1; https://eeglab.org/^80^ and available at https://github.com/slfrisby/ECoG_LASSO/. First, the CleanLine EEGlab plugin (v2.0; https://github.com/sccn/cleanline/ ^81,82^ was used to remove line noise at 60 Hz and the harmonics 120 and 180 Hz, without scanning for exact line noise frequencies and with a sliding window step size of 2 (all other parameters were set as defaults). CleanLine uses a multitaper fast Fourier transform to identify and remove sinusoidal noise and, though it removes noise less completely than traditional notch-filtering, is less disruptive to the overall frequency structure of the data. Data were then filtered using *eegfilt* (EEGLAB). The low cutoff was set to 0.5 Hz (to remove slow drifts^81^) and the high cutoff set to 300 Hz (for consistency across patients - some data were recorded at 1000 Hz using equipment that imposed a low-pass filter at 300 Hz). Channels that lay below the seizure onset zone (identified in the clinical examinations) or with poor contact (identified by visual inspection) were removed. Data were epoched between −1000 and 3000 ms relative to stimulus onset and baseline-corrected using the mean response across trials between −200 and −1 ms. Data from the nine patients recorded at 2000 Hz was then down-sampled to match the ten patients recorded at 1000 Hz by boxcar averaging pairs of neighbouring timepoints. A common average reference was applied using *reref* (EEGLAB).

Addressing artefacts in the data was a five-step process. First, the mean voltage for each channel was calculated over timepoints and trials. Any trial that contained a value more extreme than 10 standard deviations from the mean was rejected. Second, all trials were inspected individually and any that appeared to contain obvious interictal epileptiform activity, muscle activity, “electrode pop” or other artefacts were rejected. Since repeated presentations of the same stimulus were to be averaged it was important to check whether there was at least one good trial for each stimulus. One patient lacked any good trials for 18 stimuli (7 animate, 11 inanimate) and so was excluded from further analysis (and does not appear in Table 1). Another lacked any good trials for 3 stimuli (1 animate, 2 inanimate) – because of the small and relatively balanced number of missing stimuli, we included this patient. For the next steps an independent component analysis (ICA) was conducted using *runica* (EEGLAB) extracting *n* components, where *n* was 75% of the total number of good electrodes (Clarke, 2020). For the third artefact rejection step, the SASICA EEGLAB plugin (https://github.com/dnacombo/SASICA/^83^ was used to identify ICA components with weak autocorrelation, which is characteristic of muscle activity. Components identified by SASICA were inspected and, in order to minimise rejection of signal-carrying data, rejected if the channel appeared to contain only autocorrelated noise and no neural activity. Fourth, SASICA was used to highlight components with focal trial activity (i.e., activity in only a few trials, characteristic of muscle activity), “electrode pop” or other artefacts. Rather than rejecting the component across all trials, the atypical trials were identified via visual inspection and only those trials were removed. The ICA was then re-run and the process repeated until no components contained significant focal trial activity. Fifth, the possibility of microsaccade artefacts in the data was assessed^32,84–86^. The independent component activations were filtered for activity in the frequency range associated with microsaccades (20 – 190 Hz). The activations were then convolved with a saccade-related potential template and the number of microsaccades per second was calculated^87^. Since every component contained very few microsaccades (< 0.000012 microsaccades per second) and no component obviously contained more microsaccades than any other, no components were rejected based on this assessment.

Time-frequency power and phase were extracted using complex Morlet wavelet convolution^88^, implemented with *timefreq* (EEGLAB). Power values were extracted from every trial between 0 and 1650 ms in 10 ms time steps and between 4 and 200 Hz in 60 logarithmically-spaced frequency steps. A 5-cycle wavelet was used at 4 Hz, increasing to a 15-cycle wavelet at 200 Hz^32^. Power values were then averaged across repeated presentations of the same stimulus and any missing trials were interpolated with the median power value across all trials. Decibel normalisation was performed using the mean power across trials between −300 and −100 ms (calculated separately for each electrode and frequency) – a gap before stimulus onset was used to mitigate the effect of temporal leakage of trial activity into the baseline during wavelet convolution. Since phase values cannot be averaged^89^, preprocessed trials were averaged across repeated presentations of the same stimulus before phase values were extracted using the same time and frequency steps used for the extraction of power. As a comparison, preprocessed voltage was averaged over repeated presentations of the same stimulus.

The impact of preprocessing decisions on decoding results is explored in Supplementary Materials 2. In summary, decoding of semantic structure (animacy) with spectral frequency features remains unchanged after each step in this extensive and careful cleaning.

#### 3.3.3. Multivariate classification

##### 3.3.3.1. Decoding approach

Logistic regression classifiers were trained to discriminate animate from inanimate stimuli using *glmnet* in R^90^. For power and phase, frequency feature vectors were created for each stimulus at each timepoint (0 ms, 10 ms, 20 ms, …) by concatenating power or phase values for all electrodes for all frequencies in a range of interest. Voltage feature vectors were created for each stimulus at each timepoint by concatenating voltage values for all electrodes in a 50 ms window centred on the timepoint of interest. The voltage feature vectors therefore reflected activity around the timepoint of interest – but note that so did frequency feature vectors, since Morlet wavelet convolution takes activity at neighbouring timepoints into account. These vectors were provided as input to the classifiers. Each classifier was trained on vectors generated at a single timepoint for a single patient.

The classifiers used LASSO (L1) regularisation^43^, which applies a penalty that scales with the sum of the absolute values of the coefficients and thus produces solutions in which many features receive coefficients of zero. This approach was selected because it can be used to assess whether the same electrodes or frequencies are used to represent information at different timepoints. If the information used by a classifier trained at timepoint A is present at timepoint B, the classifier will perform well at timepoint B; other units, that may be in different states at the two timepoints, will receive coefficients of zero and therefore will not affect classifier performance^9^.

Classifier accuracy was assessed using 10-fold cross-validation. In each outer loop, 10 out of 100 stimuli (5 animate, 5 inanimate) were held out. The remaining 90 stimuli were used to search a range of 100 values (0.2 – 0.002, logarithmically spaced) for the regularisation parameter that resulted in the smallest mean squared error. This process was implemented using *cv.glmnet* (glmnet) with parallel model fitting implemented with foreach (https://cran.r-project.org/web/packages/foreach/index.html; all other parameters were set as defaults). A model with the best regularisation parameters was tested on the 10 stimuli in the outer loop hold-out set. The same model was tested on the same 10 stimuli at all other timepoints. The process was then repeated 10 times with different final hold-out sets. This process yielded both a main timecourse of decoding accuracy (i.e., mean accuracy, over folds, of classifiers tested on the timepoint at which they were trained) and a generalisation matrix in which the y-coordinate of a cell is the time at which the classifier was trained and the x-coordinate is the time at which the classifier was tested. Finally, a single classifier was trained at every timepoint using all 100 stimuli for training. The accuracy of these classifiers was not evaluated – they were used only for inspection of coefficients.

To assess group-level performance, the timecourses of decoding accuracy and the generalisation matrices were averaged across patients.

##### 3.3.3.2. Experimental questions

We first assessed whether power, phase and voltage each contained sufficient information to enable decoding. We therefore created both voltage feature vectors and frequency feature vectors of power and phase data (separately) using all 60 frequencies between 4 and 200 Hz and used these as input to classifiers. We compared each group-average timecourse to chance (0.5) using one-tailed, one-sample t-tests. We compared power and voltage, and phase and voltage, using paired t-tests. Probabilities were adjusted to control the false-discovery rate at α = 0.05 (Benjamini & Hochberg, 1995). We confirmed that near-identical patterns of results were obtained in the 8 patients with left-hemisphere electrodes previously analysed by Rogers et al.^9^, the 7 patients with left-hemisphere electrodes analysed for the first time in this work, and the 3 patients with right-hemisphere electrodes (Supplementary Materials 1, Figure S5). Having done so, we combined all patients into a single group for all subsequent analyses.

We next investigated whether there were differences in decoding performance between power and phase within different frequency ranges. We therefore divided the 60 frequencies into theta (4 – 7 Hz, 11 frequencies), alpha (8 - 12 Hz, 7 frequencies), beta (13 – 30 Hz, 13 frequencies), gamma (30 – 60 Hz, 10 frequencies) and high gamma (60 – 200 Hz, 19 frequencies) ranges. We constructed frequency vectors of power and phase data using only power or phase at frequencies within each range and used these as input to separate classifiers. We compared each group-average timecourse to chance and then compared each range to the timecourse of decoding using all 60 frequencies.

We then tested whether, if either power or phase data contained sufficient information to enable decoding, the power or phase code exhibited the same distributed, dynamic properties identified by Rogers et al.^9^) in both the voltage data (for the first subset of patients) and the computational hub-and-spoke model. We tested for these properties using both frequency vectors constructed using all 60 frequencies and vectors constructed using only frequencies within each range. Constant decodability had already been assessed by one-sample t-tests of each group-average timecourse against chance.

To assess local temporal generalisation we inspected the group-average generalisation matrices generated by testing each classifier at every possible timepoint. To test whether the patterns observed were statistically significant, we identified the best-performing classifier at each timepoint and then conducted paired t-tests to assess whether classifiers trained at every other possible timepoint performed equally well. Probabilities were adjusted to control the false-discovery rate at α = 0.05^91^.

To assess the shape of the generalisation window over time we computed the pairwise cosine distance between each row of the generalisation matrix, then applied agglomerative hierarchical clustering using *hclust* (native R) and cut the tree to create 10 clusters. We selected this cluster number to make these results comparable to those of Rogers et al.^9^. We averaged the timecourses of classifiers within each cluster and inspected performance over time (again, probabilities were adjusted to control the false-discovery rate). To quantify the patterns observed we took classifiers trained every 50 ms (i.e., 0 ms, 50 ms, 100 ms …; this ensured that, for classifiers trained on voltage, consecutive 50 ms windows did not overlap and were therefore independent). We calculated the area under the curve between each timecourse and a horizontal line at chance (0.5) and fitted a piecewise linear regression to the area values using the *segmented* library in R, using *selgmented* with the Bayesian Information Criterion to determine the number and location of breakpoints to produce the best model given the number of free parameters.

To assess and visualise changes in code direction we first took the classifiers trained on all data at each timepoint and then (separately for every electrode, every patient and every timepoint) calculated the mean classifier coefficient within each frequency range (or over all 60 frequencies, or over the 50 ms within each window in the case of voltage). These mean classifier coefficients were converted to one NIFTI volume per patient and per timepoint, at 2.5 mm isotropic resolution in MNI space (MNI152NLin2009cAsym), using SPM12 (https://www.fil.ion.ucl.ac.uk/spm/) implemented in MATLAB r2023b. One patient (Patient 20) was excluded from this analysis because no MNI coordinates were available for their electrodes. Other patients were missing MNI coordinates from only a subset of electrodes; we included those patients in the analysis but ignored coefficients on the electrodes with missing coordinates. We projected coefficients in each volume to the pial surface (fsaverage template^92,93^) using -*volume-to-surface-mapping* with trilinear interpolation and smoothed them on the surface at 6 mm FWHM using *–metric-smoothing* in Connectome Workbench 1.5.0. We calculated the proportion of negative coefficients that each vertex received (since animals were coded as 0 and inanimate objects as 1, a negative coefficient indicates that an increase in power is associated with increased probability that the stimulus is animate). We plotted these proportions on the pial surface using *plot_surf_stat_map* in *nilearn* 0.10.3 implemented in Python 3.9. Finally, we animated the results – ffmpeg 5.1.6 (https://ffmpeg.org/) was used to concatenate plots of coefficients at successive timepoints with a frame rate of 10 (10 times slower than real time) and motion interpolation.

## Data and code availability

We are unable to share raw data for this study because the patients did not provide informed consent to do so. However, matrices containing power, phase, and voltage features (columns) for each stimulus (rows) will be made available upon peer review and acceptance. Code is available at https://github.com/slfrisby/ECoG_LASSO/.

## CRediT statement

Saskia L. Frisby – Conceptualisation, Methodology, Formal Analysis, Data Curation, Writing – Original Draft, Writing – Review & Editing, Visualisation; Ajay D. Halai – Conceptualisation, Methodology, Formal Analysis, Writing – Review & Editing, Supervision; Christopher R. Cox – Conceptualisation, Writing – Review & Editing; Alex Clarke – Formal Analysis, Writing – Review & Editing; Akihiro Shimotake – Investigation, Data Curation, Writing – Review & Editing, Supervision; Takayuki Kikuchi – Investigation, Data Curation, Methodology; Takeharu Kuneida – Investigation, Data Curation, Methodology; Yoshiki Arakawa – Investigation, Data Curation, Methodology; Ryosuke Takahashi – Investigation; Akio Ikeda – Investigation, Data Curation, Methodology; Riki Matsumoto – Investigation, Methoology, Writing – Review & Editing; Timothy T. Rogers – Conceptualisation, Methodology, Writing – Review & Editing; Supervision; Funding Acquisition; Matthew A. Lambon Ralph - Conceptualisation, Methodology, Writing – Review & Editing; Supervision; Funding Acquisition.

## Acknowledgements

This work was supported by an MRC Career Development Award (MR/V031481/1) to A.D.H., by Grants-in-Aid for Scientific Research (KAKENHI 23KK0146, 22H02945) to R.M., and by an Advanced European Research Council (ERC) award (GAP 670428-30 BRAIN2MIND_NEUROCOMP), MRC programme grant (MR/R023883/1), and intramural funding (MC_UU_00005/18) to M.A.L.R. We would like to thank the patients who so selflessly participated in this study.

## References

1. Lambon Ralph, M.A., Jefferies, E., Patterson, K., and Rogers, T.T. (2017). The neural and computational bases of semantic cognition. Nat. Rev. Neurosci. 18, 42–55. 10.1038/nrn.2016.150.

2. Patterson, K., Nestor, P.J., and Rogers, T.T. (2007). Where do you know what you know? The representation of semantic knowledge in the human brain. Nat. Rev. Neurosci. 8, 976–987. 10.1038/nrn2277.

3. Rogers, T.T., Lambon Ralph, M.A., Garrard, P., Bozeat, S., McClelland, J.L., Hodges, J.R., and Patterson, K. (2004). Structure and deterioration of semantic memory: a neuropsychological and computational investigation. Psychol. Rev. 111, 205–235. 10.1037/0033-295X.111.1.205.

4. Başar-Eroglu, C., Strüber, D., Schürmann, M., Stadler, M., and Başar, E. (1996). Gamma-band responses in the brain: a short review of psychophysiological correlates and functional significance. Int. J. Psychophysiol. 24, 101–112. 10.1016/S0167-8760(96)00051-7.

5. Bressler, S.L. (1990). The gamma wave: a cortical information carrier? Trends Neurosci. 13, 161–162. 10.1016/0166-2236(90)90039-D.

6. Merker, B. (2013). Cortical gamma oscillations: the functional key is activation, not cognition. Neurosci. Biobehav. Rev. 37, 401–417. 10.1016/j.neubiorev.2013.01.013.

7. Nir, Y., Fisch, L., Mukamel, R., Gelbard-Sagiv, H., Arieli, A., Fried, I., and Malach, R. (2007). Coupling between Neuronal Firing Rate, Gamma LFP, and BOLD fMRI Is Related to Interneuronal Correlations. Curr. Biol. 17, 1275–1285. 10.1016/j.cub.2007.06.066.

8. Chen, Y., Shimotake, A., Matsumoto, R., Kunieda, T., Kikuchi, T., Miyamoto, S., Fukuyama, H., Takahashi, R., Ikeda, A., and Lambon Ralph, M.A. (2016). The ‘when’ and ‘where’ of semantic coding in the anterior temporal lobe: Temporal representational similarity analysis of electrocorticogram data. Cortex 79, 1–13. 10.1016/j.cortex.2016.02.015.

9. Rogers, T.T., Cox, C.R., Lu, Q., Shimotake, A., Kikuchi, T., Kunieda, T., Miyamoto, S., Takahashi, R., Ikeda, A., Matsumoto, R., et al. (2021). Evidence for a deep, distributed and dynamic code for animacy in human ventral anterior temporal cortex. eLife 10, e66276. 10.7554/eLife.66276.

10. Cox, C.R., and Rogers, T.T. (2021). Finding distributed needles in neural haystacks. J. Neurosci. 41, 1019–1032. 10.1523/JNEUROSCI.0904-20.2020.

11. Frisby, S.L., Halai, A.D., Cox, C.R., Lambon Ralph, M.A., and Rogers, T.T. (2023). Decoding semantic representations in mind and brain. Trends Cogn. Sci. 27, 258–281. 10.1016/j.tics.2022.12.006.

12. Lambon Ralph, M.A., Sage, K., Jones, R.W., and Mayberry, E.J. (2010). Coherent concepts are computed in the anterior temporal lobes. Proc. Natl. Acad. Sci. 107, 2717–2722. 10.1073/pnas.0907307107.

13. Hodges, J.R., and Patterson, K. (2007). Semantic dementia: a unique clinicopathological syndrome. Lancet Neurol. 6, 1004–1014. 10.1016/S1474-4422(07)70266-1.

14. Devlin, J.T., Russell, R.P., Davis, M.H., Price, C.J., Wilson, J., Moss, H.E., Matthews, P.M., and Tyler, L.K. (2000). Susceptibility-induced loss of signal: Comparing PET and fMRI on a semantic task. NeuroImage 11, 589–600. 10.1006/nimg.2000.0595.

15. Binney, R.J., Embleton, K.V., Jefferies, E., Parker, G.J.M., and Lambon Ralph, M.A. (2010). The ventral and inferolateral aspects of the anterior temporal lobe are crucial in semantic memory: Evidence from a novel direct comparison of distortion-corrected fMRI, rTMS, and semantic dementia. Cereb. Cortex 20, 2728–2738. 10.1093/cercor/bhq019.

16. Halai, A.D., Welbourne, S.R., Embleton, K., and Parkes, L.M. (2014). A comparison of dual gradient-echo and spin-echo fMRI of the inferior temporal lobe. Hum. Brain Mapp. 35, 4118–4128. 10.1002/hbm.22463.

17. Visser, M., Jefferies, E., and Lambon Ralph, M.A. (2010). Semantic processing in the anterior temporal lobes: a meta-analysis of the functional neuroimaging literature. J. Cogn. Neurosci. 22, 1083–1094. 10.1162/jocn.2009.21309.

18. Mollo, G., Cornelissen, P.L., Millman, R.E., Ellis, A.W., and Jefferies, E. (2017). Oscillatory dynamics supporting semantic cognition: MEG evidence for the contribution of the anterior temporal lobe hub and modality-specific spokes. PLOS ONE 12, e0169269. 10.1371/journal.pone.0169269.

19. Chen, L., Lambon Ralph, M.A., and Rogers, T.T. (2017). A unified model of human semantic knowledge and its disorders. Nat. Hum. Behav. 1, 0039. 10.1038/s41562-016-0039.

20. Pobric, G., Jefferies, E., and Lambon Ralph, M.A. (2007). Anterior temporal lobes mediate semantic representation: Mimicking semantic dementia by using rTMS in normal participants. Proc. Natl. Acad. Sci. 104, 20137–20141. 10.1073/pnas.0707383104.

21. Pobric, G., Jefferies, E., and Lambon Ralph, M.A. (2010). Amodal semantic representations depend on both anterior temporal lobes: Evidence from repetitive transcranial magnetic stimulation. Neuropsychologia 48, 1336–1342. 10.1016/j.neuropsychologia.2009.12.036.

22. Pobric, G., Jefferies, E., and Lambon Ralph, M.A. (2010). Category-specific versus category-general semantic impairment induced by transcranial magnetic stimulation. Curr. Biol. 20, 964–968. 10.1016/j.cub.2010.03.070.

23. Lüders, H., Lesser, R.P., Hahn, J., Dinner, D.S., Morris, H.H., Wylie, E., and Godoy, J. (1991). Basal temporal language area. Brain 114, 743–754. 10.1093/brain/114.2.743.

24. Matoba, K., Matsumoto, R., Shimotake, A., Nakae, T., Imamura, H., Togo, M., Yamao, Y., Usami, K., Kikuchi, T., Yoshida, K., et al. (2024). Basal temporal language area revisited in Japanese language with a language function density map. Cereb. Cortex 34, bhae218. 10.1093/cercor/bhae218.

25. Halgren, E., Wang, C., Schomer, D.L., Knake, S., Marinkovic, K., Wu, J., and Ulbert, I. (2006). Processing stages underlying word recognition in the anteroventral temporal lobe. NeuroImage 30, 1401–1413. 10.1016/j.neuroimage.2005.10.053.

26. Nobre, A., and McCarthy, G. (1995). Language-related field potentials in the anterior-medial temporal lobe: II. Effects of word type and semantic priming. J. Neurosci. 15, 1090–1098. 10.1523/JNEUROSCI.15-02-01090.1995.

27. Sato, N., Matsumoto, R., Shimotake, A., Matsuhashi, M., Otani, M., Kikuchi, T., Kunieda, T., Mizuhara, H., Miyamoto, S., Takahashi, R., et al. (2021). Frequency-dependent cortical interactions during semantic processing: an electrocorticogram cross-spectrum analysis using a semantic space model. Cereb. Cortex 31, 4329–4339. 10.1093/cercor/bhab089.

28. Shimotake, A., Matsumoto, R., Ueno, T., Kunieda, T., Saito, S., Hoffman, P., Kikuchi, T., Fukuyama, H., Miyamoto, S., Takahashi, R., et al. (2015). Direct exploration of the role of the ventral anterior temporal lobe in semantic memory: cortical stimulation and local field potential evidence from subdural grid electrodes. Cereb. Cortex 25, 3802–3817. 10.1093/cercor/bhu262.

29. Cox, C.R., Rogers, T.T., Shimotake, A., Kikuchi, T., Kunieda, T., Miyamoto, S., Takahashi, R., Matsumoto, R., Ikeda, A., and Lambon Ralph, M.A. (2024). Representational similarity learning reveals a graded multidimensional semantic space in the human anterior temporal cortex. Imaging Neurosci. 2, 1–22. 10.1162/imag_a_00093.

30. Liu, H., Agam, Y., Madsen, J.R., and Kreiman, G. (2009). Timing, timing, timing: Fast decoding of object information from intracranial field potentials in human visual cortex. Neuron 62, 281–290. 10.1016/j.neuron.2009.02.025.

31. Nagata, K., Kunii, N., Shimada, S., Fujitani, S., Takasago, M., and Saito, N. (2022). Spatiotemporal target selection for intracranial neural decoding of abstract and concrete semantics. Cereb. Cortex 32, 5544–5554. 10.1093/cercor/bhac034.

32. Clarke, A. (2020). Dynamic activity patterns in the anterior temporal lobe represents object semantics. Cogn. Neurosci. 11, 111–121. 10.1080/17588928.2020.1742678.

33. Rupp, K., Roos, M., Milsap, G., Caceres, C., Ratto, C., Chevillet, M., Crone, N.E., and Wolmetz, M. (2017). Semantic attributes are encoded in human electrocorticographic signals during visual object recognition. NeuroImage 148, 318–329. 10.1016/j.neuroimage.2016.12.074.

34. Wang, W., Degenhart, A.D., Sudre, G.P., Pomerleau, D.A., and Tyler-Kabara, E.C. (2011). Decoding semantic information from human electrocorticographic (ECoG) signals. Annu. Int. Conf. IEEE Eng. Med. Biol. Soc. IEEE Eng. Med. Biol. Soc. Annu. Int. Conf. 2011, 6294–6298. 10.1109/IEMBS.2011.6091553.

35. Canolty, R.T., and Knight, R.T. (2010). The functional role of cross-frequency coupling. Trends Cogn. Sci. 14, 506–515. 10.1016/j.tics.2010.09.001.

36. Lachaux, J.-P., Axmacher, N., Mormann, F., Halgren, E., and Crone, N.E. (2012). High-frequency neural activity and human cognition: Past, present and possible future of intracranial EEG research. Prog. Neurobiol. 98, 279–301. 10.1016/j.pneurobio.2012.06.008.

37. Parvizi, J., and Kastner, S. (2018). Promises and limitations of human intracranial electroencephalography. Nat. Neurosci. 21, 474–483. 10.1038/s41593-018-0108-2.

38. Carlson, T., Tovar, D.A., Alink, A., and Kriegeskorte, N. (2013). Representational dynamics of object vision: The first 1000 ms. J. Vis. 13, 1–1. 10.1167/13.10.1.

39. Cichy, R.M., Pantazis, D., and Oliva, A. (2014). Resolving human object recognition in space and time. Nat. Neurosci. 17, 455–462. 10.1038/nn.3635.

40. Contini, E.W., Wardle, S.G., and Carlson, T.A. (2017). Decoding the time-course of object recognition in the human brain: From visual features to categorical decisions. Neuropsychologia 105, 165–176. 10.1016/j.neuropsychologia.2017.02.013.

41. King, J.-R., and Dehaene, S. (2014). Characterizing the dynamics of mental representations: the temporal generalization method. Trends Cogn. Sci. 18, 203–210. 10.1016/j.tics.2014.01.002.

42. Dilkina, K., and Lambon Ralph, M.A. (2012). Conceptual structure within and between modalities. Front. Hum. Neurosci. 6. 10.3389/fnhum.2012.00333.

43. Tibshirani, R. (1996). Regression shrinkage and selection via the lasso. J. R. Stat. Soc. Ser. B Methodol. 58, 267–288. 10.1111/j.2517-6161.1996.tb02080.x.

44. Hermes, D., Miller, K.J., Vansteensel, M.J., Edwards, E., Ferrier, C.H., Bleichner, M.G., van Rijen, P.C., Aarnoutse, E.J., and Ramsey, N.F. (2014). Cortical theta wanes for language. NeuroImage 85, 738–748. 10.1016/j.neuroimage.2013.07.029.

45. Clarke, A., Devereux, B.J., and Tyler, L.K. (2018). Oscillatory dynamics of perceptual to conceptual transformations in the ventral visual pathway. J. Cogn. Neurosci. 30, 1590–1605. 10.1162/jocn_a_01325.

46. Abel, T.J., Rhone, A.E., Nourski, K.V., Kawasaki, H., Oya, H., Griffiths, T.D., Howard, M.A., and Tranel, D. (2015). Direct physiologic evidence of a heteromodal convergence region for proper naming in human left anterior temporal lobe. J. Neurosci. 35, 1513–1520. 10.1523/JNEUROSCI.3387-14.2015.

47. Arya, R. (2019). Similarity of spatiotemporal dynamics of language-related ECoG high-gamma modulation in Japanese and English speakers. Clin. Neurophysiol. Off. J. Int. Fed. Clin. Neurophysiol. 130, 1403–1404. 10.1016/j.clinph.2019.05.006.

48. Cervenka, M.C., Corines, J., Boatman-Reich, D.F., Eloyan, A., Sheng, X., Franaszczuk, P.J., and Crone, N.E. (2013). Electrocorticographic functional mapping identifies human cortex critical for auditory and visual naming. NeuroImage 69, 267–276. 10.1016/j.neuroimage.2012.12.037.

49. Chan, A.M., Baker, J.M., Eskandar, E., Schomer, D., Ulbert, I., Marinkovic, K., Cash, S.S., and Halgren, E. (2011). First-pass selectivity for semantic categories in human anteroventral temporal lobe. J. Neurosci. 31, 18119–18129. 10.1523/JNEUROSCI.3122-11.2011.

50. Crone, N.E., Boatman, D., Gordon, B., and Hao, L. (2001). Induced electrocorticographic gamma activity during auditory perception. Clin. Neurophysiol. 112, 565–582. 10.1016/S1388-2457(00)00545-9.

51. Crone, N.E., and Hao, L. (2002). Functional dynamics of spoken and signed word production: A case study using electrocorticographic spectral analysis. Aphasiology 16, 903–927. 10.1080/02687030244000383.

52. Edwards, E., Nagarajan, S.S., Dalal, S.S., Canolty, R.T., Kirsch, H.E., Barbaro, N.M., and Knight, R.T. (2010). Spatiotemporal imaging of cortical activation during verb generation and picture naming. NeuroImage 50, 291–301. 10.1016/j.neuroimage.2009.12.035.

53. Forseth, K.J., Kadipasaoglu, C.M., Conner, C.R., Hickok, G., Knight, R.T., and Tandon, N. (2018). A lexical semantic hub for heteromodal naming in middle fusiform gyrus. Brain 141, 2112–2126. 10.1093/brain/awy120.

54. Kojima, K., Brown, E.C., Matsuzaki, N., Rothermel, R., Fuerst, D., Shah, A., Mittal, S., Sood, S., and Asano, E. (2013). Gamma activity modulated by picture and auditory naming tasks: Intracranial recording in patients with focal epilepsy. Clin. Neurophysiol. 124, 1737–1744. 10.1016/j.clinph.2013.01.030.

55. Nakai, Y., Jeong, J., Brown, E.C., Rothermel, R., Kojima, K., Kambara, T., Shah, A., Mittal, S., Sood, S., and Asano, E. (2017). Three- and four-dimensional mapping of speech and language in patients with epilepsy. Brain 140, 1351–1370. 10.1093/brain/awx051.

56. Nakai, Y., Sugiura, A., Brown, E.C., Sonoda, M., Jeong, J., Rothermel, R., Luat, A.F., Sood, S., and Asano, E. (2019). Four-dimensional functional cortical maps of visual and auditory language: Intracranial recording. Epilepsia 60, 255–267. 10.1111/epi.14648.

57. Snyder, K.M., Forseth, K.J., Donos, C., Rollo, P.S., Fischer-Baum, S., Breier, J., and Tandon, N. (2023). Critical role of the ventral temporal lobe in naming. Epilepsia 64, 1200–1213. 10.1111/epi.17555.

58. Tanji, K. (2005). High-frequency -band activity in the basal temporal cortex during picture-naming and lexical-decision tasks. J. Neurosci. 25, 3287–3293. 10.1523/JNEUROSCI.4948-04.2005.

59. Jones, M.N., Willits, J., and Dennis, S. (2015). In Models of Semantic Memory, J. R. Busemeyer, Z. Wang, J. T. Townsend, and A. Eidels, eds. (Oxford University Press). 10.1093/oxfordhb/9780199957996.013.11.

60. Yuste, R. (2015). From the neuron doctrine to neural networks. Nat. Rev. Neurosci. 16, 487–497. 10.1038/nrn3962.

61. Drane, D.L., Ojemann, G.A., Aylward, E., Ojemann, J.G., Johnson, L.C., Silbergeld, D.L., Miller, J.W., and Tranel, D. (2008). Category-specific naming and recognition deficits in temporal lobe epilepsy surgical patients. Neuropsychologia 46, 1242–1255. 10.1016/j.neuropsychologia.2007.11.034.

62. Hamberger, M.J. (2007). Cortical language mapping in epilepsy: A critical review. Neuropsychol. Rev. 17, 477–489. 10.1007/s11065-007-9046-6.

63. Henry, T.R., Buchtel, H.A., Koeppe, R.A., Pennell, P.B., Kluin, K.J., and Minoshima, S. (1998). Absence of normal activation of the left anterior fusiform gyrus during naming in left temporal lobe epilepsy. Neurology 50, 787–790. 10.1212/WNL.50.3.787.

64. Watrous, A.J., Fell, J., Ekstrom, A.D., and Axmacher, N. (2015). More than spikes: common oscillatory mechanisms for content specific neural representations during perception and memory. Curr. Opin. Neurobiol. 31, 33–39. 10.1016/j.conb.2014.07.024.

65. Sauseng, P., and Klimesch, W. (2008). What does phase information of oscillatory brain activity tell us about cognitive processes? Neurosci. Biobehav. Rev. 32, 1001–1013. 10.1016/j.neubiorev.2008.03.014.

66. Stryker, M.P. (1989). Is grandmother an oscillation? Nature 338, 297–298. 10.1038/338297a0.

67. Tallon-Baudry, C., and Bertrand, O. (1999). Oscillatory gamma activity in humans and its role in object representation. Trends Cogn. Sci. 3, 151–162. 10.1016/S1364-6613(99)01299-1.

68. von der Malsburg, C. (1995). Binding in models of perception and brain function. Curr. Opin. Neurobiol. 5, 520–526. 10.1016/0959-4388(95)80014-X.

69. Solomon, E.A., Lega, B.C., Sperling, M.R., and Kahana, M.J. (2019). Hippocampal theta codes for distances in semantic and temporal spaces. 10.

70. Spiers, H.J. (2020). The hippocampal cognitive map: One space or many? Trends Cogn. Sci. 24, 168–170. 10.1016/j.tics.2019.12.013.

71. Halgren, E., Kaestner, E., Marinkovic, K., Cash, S.S., Wang, C., Schomer, D.L., Madsen, J.R., and Ulbert, I. (2015). Laminar profile of spontaneous and evoked theta: Rhythmic modulation of cortical processing during word integration. Neuropsychologia 76, 108–124. 10.1016/j.neuropsychologia.2015.03.021.

72. Jackson, R.L. (2021). The neural correlates of semantic control revisited. NeuroImage 224, 117444. 10.1016/j.neuroimage.2020.117444.

73. Klimesch, W. (2012). Alpha-band oscillations, attention, and controlled access to stored information. Trends Cogn. Sci. 16, 606–617. 10.1016/j.tics.2012.10.007.

74. Marko, M., Cimrová, B., and Riečanský, I. (2019). Neural theta oscillations support semantic memory retrieval. Sci. Rep. 9, 17667. 10.1038/s41598-019-53813-y.

75. Cox, C.R., Seidenberg, M.S., and Rogers, T.T. (2015). Connecting functional brain imaging and Parallel Distributed Processing. Lang. Cogn. Neurosci. 30, 380–394. 10.1080/23273798.2014.994010.

76. Morrison, C.M., Chappell, T.D., and Ellis, A.W. (1997). Age of acquisition norms for a large set of object names and their relation to adult estimates and other variables. Q. J. Exp. Psychol. Sect. A 50, 528–559. 10.1080/027249897392017.

77. Barry, C., Morrison, C.M., and Ellis, A.W. (1997). Naming the Snodgrass and Vanderwart pictures: Effects of age of acquisition, frequency, and name agreement. Q. J. Exp. Psychol. Sect. A 50, 560–585. 10.1080/783663595.

78. Jenkinson, M., Beckmann, C.F., Behrens, T.E.J., Woolrich, M.W., and Smith, S.M. (2012). FSL. NeuroImage 62, 782–790. 10.1016/j.neuroimage.2011.09.015.

79. Smith, S.M., Jenkinson, M., Woolrich, M.W., Beckmann, C.F., Behrens, T.E.J., Johansen-Berg, H., Bannister, P.R., De Luca, M., Drobnjak, I., Flitney, D.E., et al. (2004). Advances in functional and structural MR image analysis and implementation as FSL. NeuroImage 23, S208–S219. 10.1016/j.neuroimage.2004.07.051.

80. Delorme, A., and Makeig, S. (2004). EEGLAB: an open source toolbox for analysis of single-trial EEG dynamics including independent component analysis. J. Neurosci. Methods 134, 9–21. 10.1016/j.jneumeth.2003.10.009.

81. Delorme, A. (2023). EEG is better left alone. Sci. Rep. 13, 2372. 10.1038/s41598-023-27528-0.

82. Mitra, P., and Bokil, H. (2007). Observed Brain Dynamics (Oxford University Press) 10.1093/acprof:oso/9780195178081.001.0001.

83. Chaumon, M., Bishop, D.V.M., and Busch, N.A. (2015). A practical guide to the selection of independent components of the electroencephalogram for artifact correction. J. Neurosci. Methods 250, 47–63. 10.1016/j.jneumeth.2015.02.025.

84. Jerbi, K., Freyermuth, S., Dalal, S., Kahane, P., Bertrand, O., Berthoz, A., and Lachaux, J.-P. (2009). Saccade related gamma-band activity in intracerebral EEG: Dissociating neural from ocular muscle activity. Brain Topogr. 22, 18–23. 10.1007/s10548-009-0078-5.

85. Kovach, C.K., Tsuchiya, N., Kawasaki, H., Oya, H., Howard, M.A., and Adolphs, R. (2011). Manifestation of ocular-muscle EMG contamination in human intracranial recordings. NeuroImage 54, 213–233. 10.1016/j.neuroimage.2010.08.002.

86. Yuval-Greenberg, S., Tomer, O., Keren, A.S., Nelken, I., and Deouell, L.Y. (2008). Transient induced gamma-band response in eeg as a manifestation of miniature saccades. Neuron 58, 429–441. 10.1016/j.neuron.2008.03.027.

87. Craddock, M., Martinovic, J., and Müller, M.M. (2016). Accounting for microsaccadic artifacts in the EEG using independent component analysis and beamforming. Psychophysiology 53, 553–565. 10.1111/psyp.12593.

88. Bertrand, O., Bohorquez, J., and Pernier, J. (1994). Time-frequency digital filtering based on an invertible wavelet transform: an application to evoked potentials. IEEE Trans. Biomed. Eng. 41, 77–88. 10.1109/10.277274.

89. Cohen, M.X. (2014). Analyzing Neural Time Series Data: Theory and Practice (MIT Press).

90. Friedman, J., Hastie, T., and Tibshirani, R. (2010). Regularization paths for generalized linear models via coordinate descent. J. Stat. Softw. 33. 10.18637/jss.v033.i01.

91. Benjamini, Y., and Hochberg, Y. (1995). Controlling the false discovery rate: A practical and powerful approach to multiple testing. J. R. Stat. Soc. Ser. B Stat. Methodol. 57, 289–300.

92. Dale, A.M., Fischl, B., and Sereno, M.I. (1999). Cortical surface-based analysis: I. Segmentation and surface reconstruction. NeuroImage 9, 179–194. 10.1006/nimg.1998.0395.

93. Fischl, B., Sereno, M.I., and Dale, A.M. (1999). Cortical surface-based analysis: II. Inflation, flattening, and a surface-based coordinate system. NeuroImage 9, 195–207. 10.1006/nimg.1998.0396.

