## Supplementary materials 1 for "All spectral frequencies of neural activity reveal semantic representation in the human anterior ventral temporal cortex"

### **Supplementary Materials S1: Additional analyses**

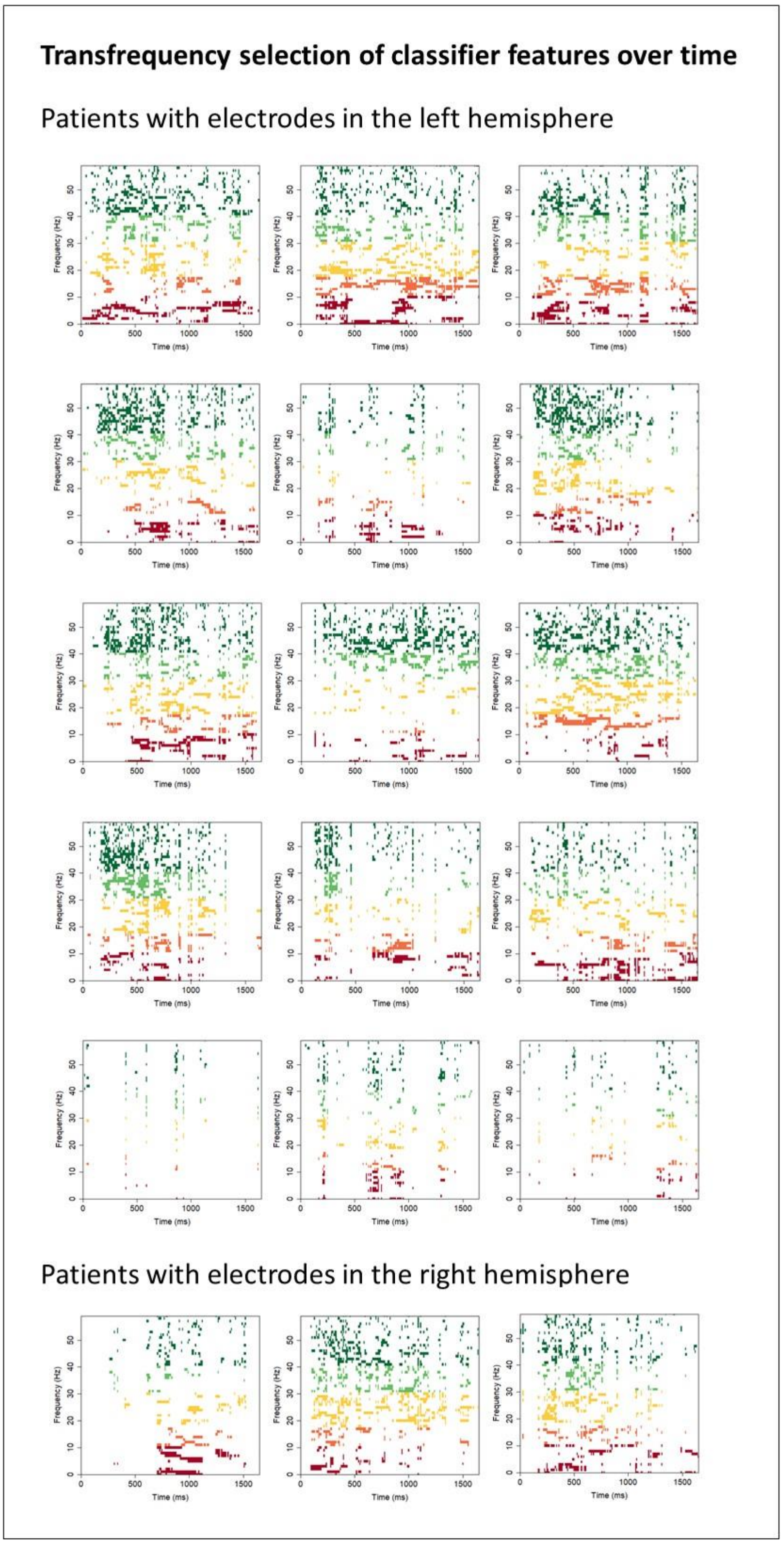


Figure S1: Feature selection of each frequency at each timepoint (averaged across electrodes). Classifiers were trained on power frequency features for all frequencies between 4 and 200 Hz. Features with a nonzero coefficient are shown in colour - theta (4 – 7 Hz, red), alpha (8 – 12 Hz, orange), beta (13 – 30 Hz, yellow), gamma (30 – 60 Hz, light green) and high gamma (60 – 200 Hz, dark green). Each plot shows selection for a single patient.

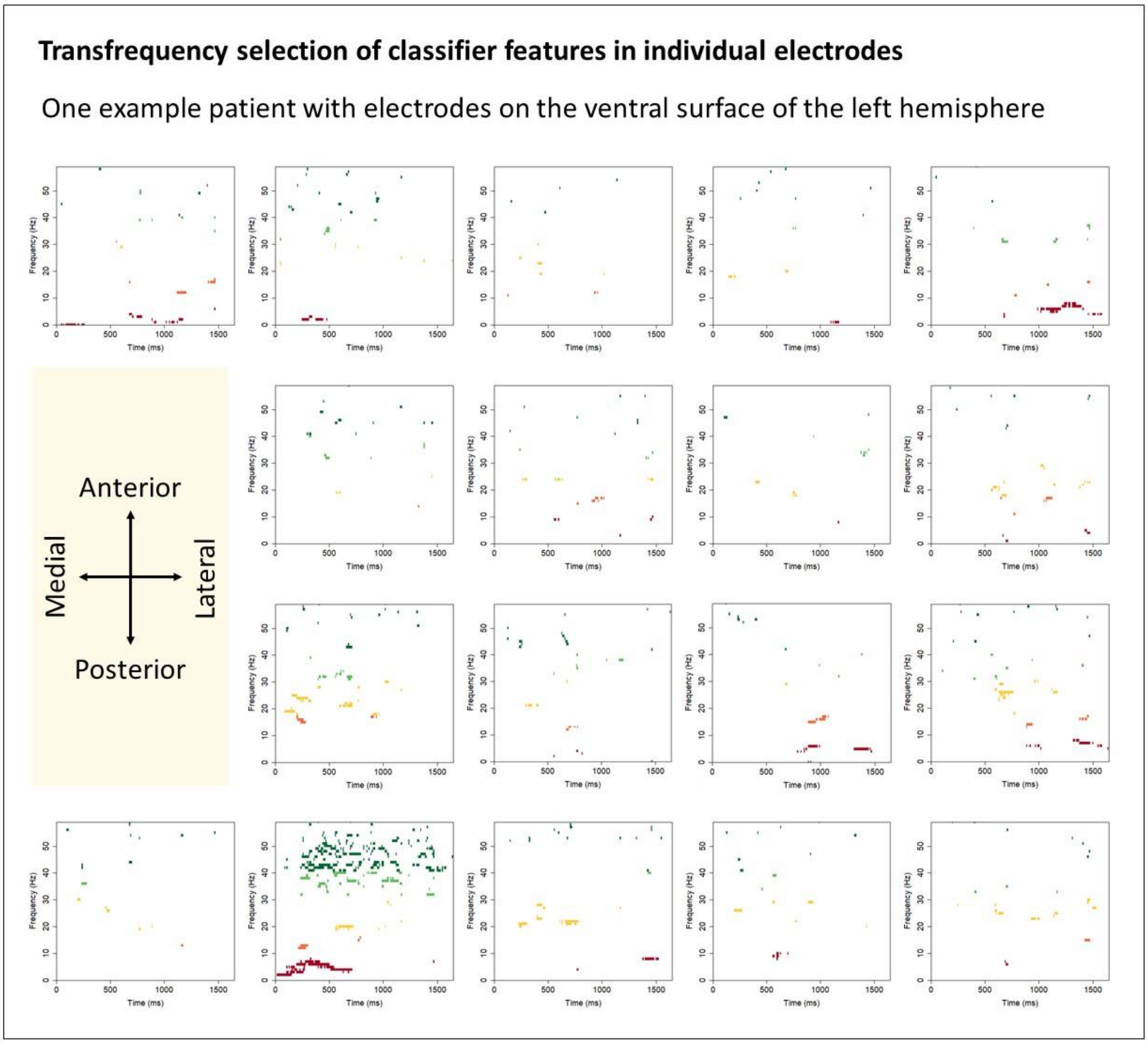


Figure S2: Feature selection in each electrode for a single patient. Classifiers were trained on power frequency features for all frequencies between 4 and 200 Hz. Features with a nonzero coefficient are shown in colour - theta (4 – 7 Hz, red), alpha (8 – 12 Hz, orange), beta (13 – 30 Hz, yellow), gamma (30 – 60 Hz, light green) and high gamma (60 – 200 Hz, dark green). Each plot shows coefficients for a single electrode. The arrangement of the plots illustrates the layout of the electrodes on the ventral surface of the left hemisphere of this patient's brain.

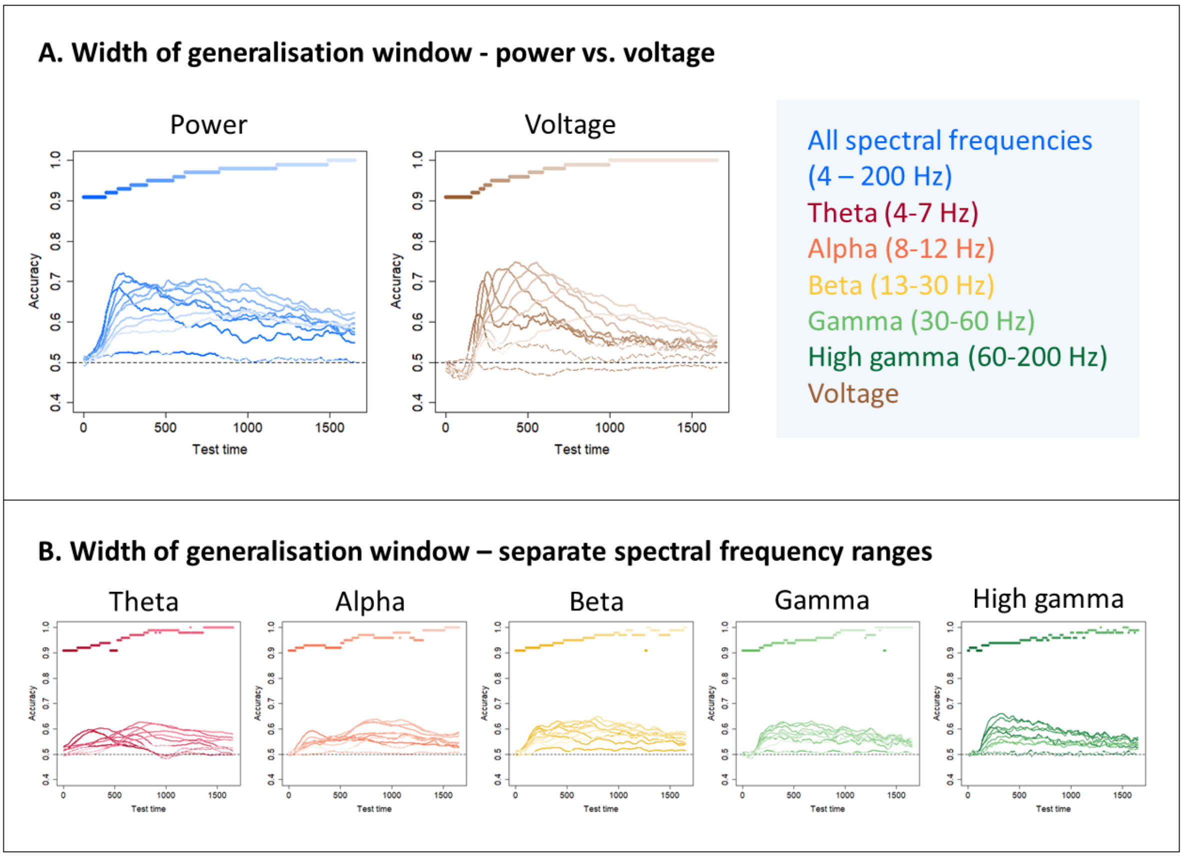


Figure S3: Width of generalisation window for all frequency ranges. Classifiers are grouped into 10 clusters via agglomerative hierarchical clustering (see Methods). The "timecourses" show the mean hold-out accuracy for classifiers within each cluster at each timepoint. Lines are solid where there is a significant difference between classifier accuracy and chance (0.5, one-sample t-test with probabilities adjusted to control the false-discovery rate at $\alpha = 0.05$) and dashed where there is no significant difference. Coloured bars show the grouped timepoints in each cluster. (A) Results for classifiers trained on power frequency features for all frequencies between 4 and 200 Hz (shades of blue) or on voltage features (shades of brown). (B) Results for classifiers trained on power frequency features from a single range - theta (4 – 7 Hz, shades of red), alpha (8 - 12 Hz, shades of orange), beta (13 – 30 Hz, shades of yellow), gamma (30 – 60 Hz, shades of light green) and high gamma (60 – 200 Hz, shades of dark green).

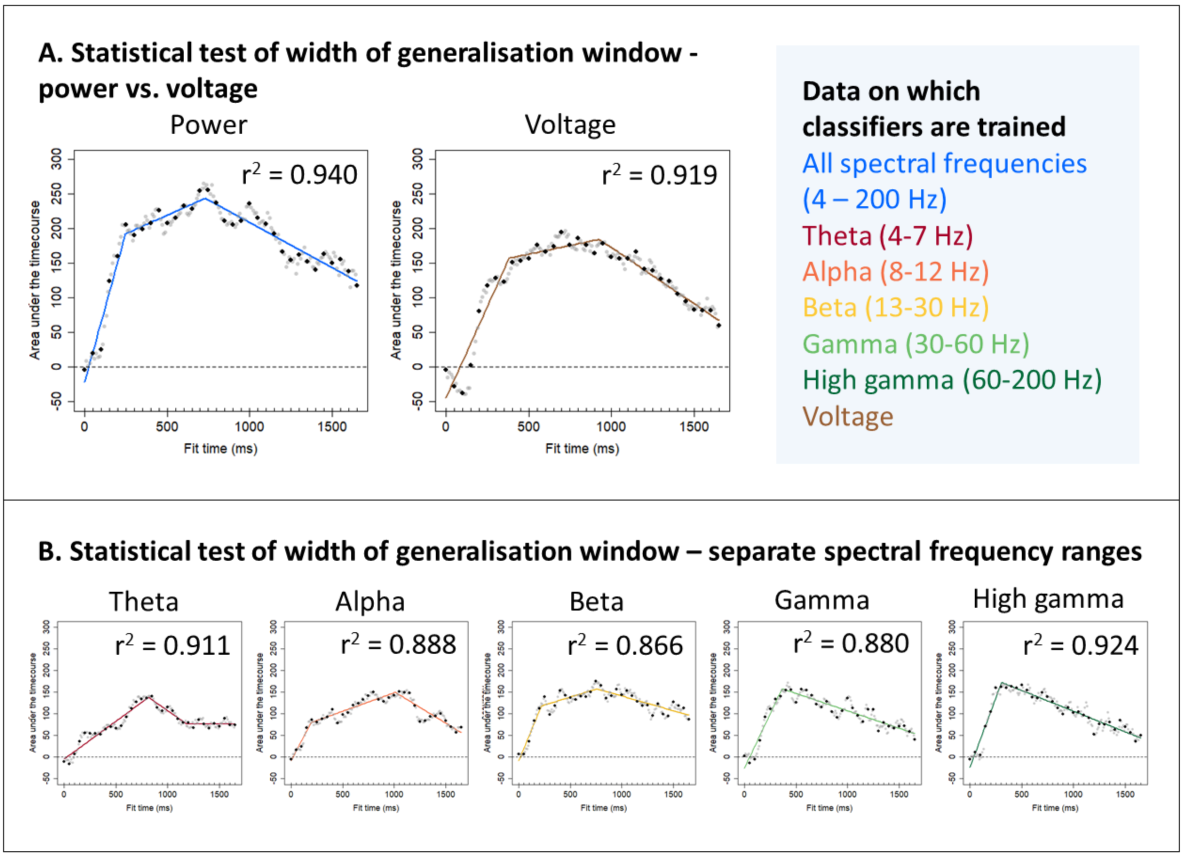


Figure S4: Statistical test of width of generalisation window. Plots show area under the curve between the timecourse of hold-out accuracy for each classifier and a horizontal line at chance (0.5). Lines show a piecewise linear model fit to timepoints 50 ms apart (0 ms, 50 ms, 100 ms, …; black dots). For each model segment, a positive gradient indicates that the generalisation window is widening. Fitting a model to timepoints 50 ms apart ensures that voltage feature vectors from neighbouring timepoints do not contain overlapping data points and that power results are comparable with voltage results. Timepoints which were not used to fit the model are shown in grey. (A) Results for classifiers trained on power frequency features for all frequencies between 4 and 200 Hz (blue) or on voltage features (brown). (B) Results for classifiers trained on power frequency features from a single range - theta (4 – 7 Hz, red), alpha (8 - 12 Hz, orange), beta (13 – 30 Hz, yellow), gamma (30 – 60 Hz, light green) and high gamma (60 – 200 Hz, dark green).

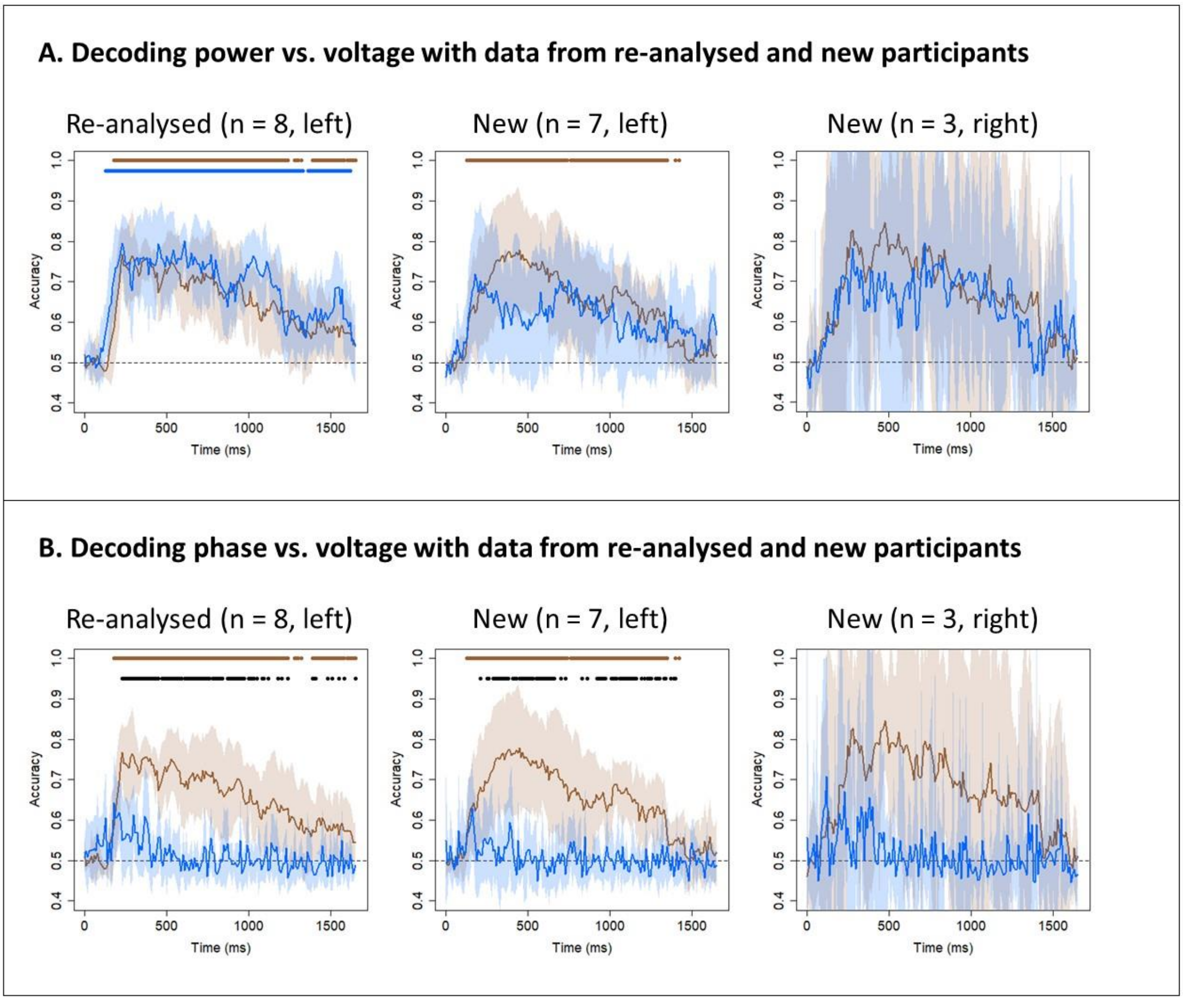


Figure S5: Decoding subsamples of patients. (A) Mean and 95% confidence interval of the hold-out accuracy for classifiers trained on power frequency features for all frequencies between 4 and 200 Hz (blue) or on voltage features (brown) from three subsamples of patients – the re-analysed 8 patients (originally analysed by Rogers et al., 2021), the 7 left-hemisphere patients analysed for the first time in this work, and the 3 right-hemisphere patients analysed for the first time in this work. Coloured dots indicate a significant difference between classifier accuracy and chance (0.5, one-sample t-test with probabilities adjusted to control the false-discovery rate at $\alpha = 0.05$). Black dots indicate a significant difference between accuracies at a given timepoint (paired t-tests with probabilities adjusted to control the false-discovery rate at $\alpha = 0.05$). (B) Mean and 95% confidence interval of the hold-out accuracy for classifiers trained on phase frequency features (blue) or on voltage features (brown). Coloured dots indicate a significant difference between classifier accuracy and chance; black dots indicate a significant difference between accuracies.

### References

Rogers, T. T., Cox, C. R., Lu, Q., Shimotake, A., Kikuchi, T., Kunieda, T., Miyamoto, S., Takahashi, R., Ikeda, A., Matsumoto, R., & Lambon Ralph, M. A. (2021). Evidence for a deep, distributed and dynamic code for animacy in human ventral anterior temporal cortex. *eLife*, *10*, e66276. https://doi.org/10.7554/eLife.66276
