## Supplementary materials 2 for "All spectral frequencies of neural activity reveal semantic representation in the human anterior ventral temporal cortex"

**Supplementary Materials S2: The impact of preprocessing on decoding accuracy**

### 1. Introduction

We formally checked two possible technical issues within the preprocessing pipeline. The first was that preprocessing may remove signal as well as noise (de Cheveigné, 2023; Delorme, 2023) and so may harm decoding performance. The second was that, since electrooculogram (EOG) data were available for only 5 patients, it was not possible to identify and correct for macrosaccades directly. This factor is important to examine given that saccadic activity may differ between conditions – the silhouettes of living things tend to be more similar to each other than the silhouettes of nonliving things are, which means that greater attention to visual detail is needed to identify living things (Humphreys & Forde, 2001). There is a notional possibility, therefore, that if microsaccades can influence the cortical electrode data, then this difference in eye movement might be enough to drive or contribute to decoding performance.

### 2. Methods

#### 2.1. Preprocessing – ECoG

Preprocessing followed the pipeline described in the main text. To summarise, we implemented the following steps:

1. Filtering (CleanLine to remove line noise at 60 Hz and the harmonics 120 and 180 Hz, then filtering with low cutoff 0.5 Hz and high cutoff 300 Hz)
2. Rejection of channels below the seizure onset zone or with poor contact
3. Application of a common average reference
4. Rejection of trials that contained obvious interictal epileptiform activity, muscle activity, "electrode pop" or other artefacts

Raw data and data after each preprocessing step (to illustrate – data after filtering, data after filtering *and* channel rejection, etc.) were epoched between -1000 and 3000 ms relative to stimulus onset and baseline-corrected using the mean response across trials between -200 and -1 ms. Data from the 9 patients recorded at 2000 Hz was then downsampled to match the 10 patients recorded at 1000 Hz by boxcar averaging pairs of neighbouring timepoints. For the 5 patients for whom we had EOG data, we also epoched, baseline-corrected and downsampled the EOG data (4 patients had a single EOG channel and the other had 2). No other preprocessing was performed on the EOG data.

Time-frequency power and phase were extracted from data at each preprocessing step using complex Morlet wavelet convolution with the same parameters used in the main analysis. Power was averaged across repeated presentations of the same stimulus, any missing trials were interpolated, and decibel normalisation was performed using the same parameters used in the main analysis. For phase, preprocessed trials were averaged across repeated presentations of the same stimulus before phase values were extracted. Power and phase were extracted from EOG data in the same way. As a comparison, preprocessed voltage was averaged over repeated presentations of the same stimulus.

### 2.2. Multivariate classification

#### 2.2.1. Decoding approach

The decoding approach was identical to that described in the main text.

#### 2.2.2. Experimental questions

To address the first query (whether preprocessing may remove signal as well as noise), we created frequency feature vectors (including all 60 frequencies) and voltage feature vectors from raw data and data after each step of preprocessing. We used these as input to classifiers and compared each group-average timecourse to chance (0.5) using one-tailed, one-sample t-tests with false discovery rate correction as described in the main text. Where there was a visible difference in accuracy before and after a preprocessing step, we compared the timecourse before that step to the timecourse after that step using paired t-tests with false discovery rate correction as described in the main text.

To address the second query (whether saccadic activity may drive decoding performance), we created frequency feature vectors and voltage feature vectors from power, phase and voltage values extracted from EOG data. We hypothesised that eye movements may affect decoding results from different frequency ranges differently and so we created frequency vectors for all 60 frequencies and then for each range individually. We averaged the timecourses over the 5 participants with EOG data available and used paired t-tests with false discovery rate correction to compare the group-average timecourse of decoding EOG power, phase, or voltage to the average timecourse of decoding ECoG power, phase, or voltage.

### 3. Results

Figure S7 shows the decoding profile of classifiers fitted to power, phase and voltage data – first to raw data, then to data after each preprocessing step (filtering, channel rejection, rereferencing and trial rejection). The decoding profile of time-frequency power and phase hardly changed when preprocessing steps were applied (a slight decrease in accuracy pushed phase decoding below the threshold for significance at some timepoints). By contrast, while voltage was significantly decodable throughout the time window of interest, the shape of the decoding profile changed after filtering – decoding accuracy reached the same maximum but declined more steeply from that point. After around 500 ms, accuracy for classifiers fitted to filtered data was significantly poorer than accuracy for classifiers fitted to raw data.

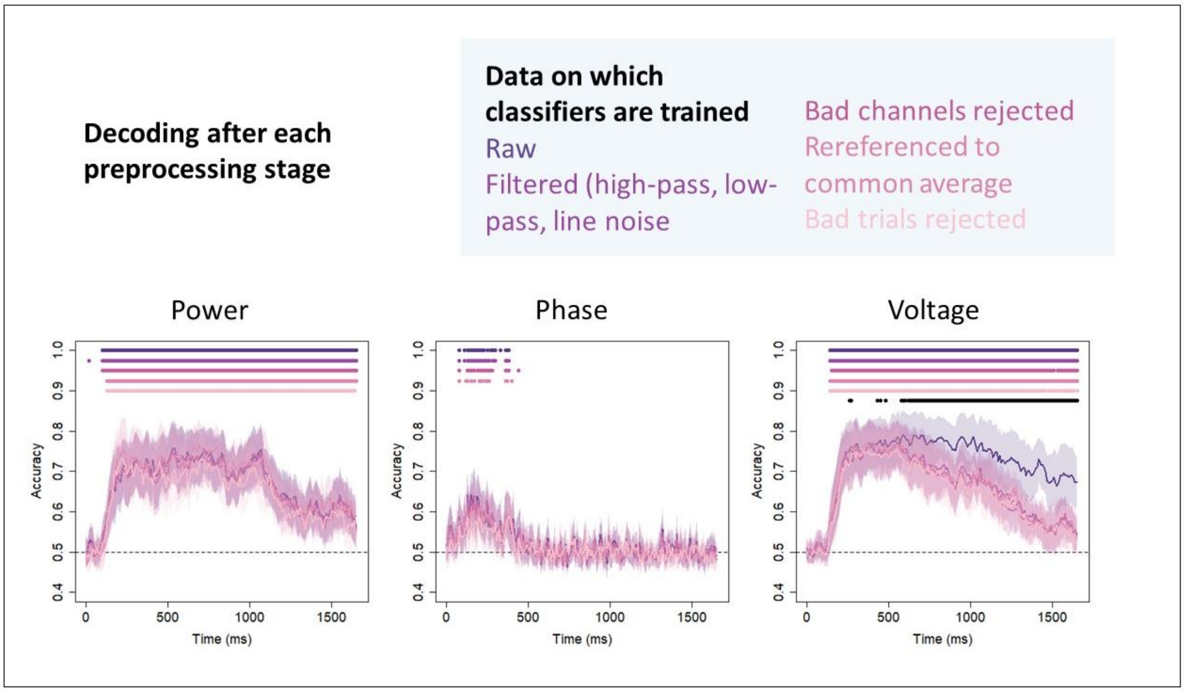


Figure S7: Decoding during preprocessing. Mean and 95% confidence interval of the hold-out accuracy for classifiers trained on power or phase frequency features, including all 60 frequencies between 4 and 200 Hz, or on voltage features. Classifiers are trained after each preprocessing step – none (raw data; dark purple), after filtering (light purple), after bad channels were rejected (dark pink), after common average referencing (medium pink), and after bad trials were rejected (pale pink). Coloured dots indicate a significant difference between classifier accuracy and chance (0.5, one-sample t-test with probabilities adjusted to control the false-discovery rate at $\alpha = 0.05$). Black dots indicate a significant difference between accuracy for classifiers trained on voltage extracted from raw data (dark purple) and classifiers trained on voltage extracted from data after filtering (light purple) at a given timepoint (paired t-tests with probabilities adjusted to control the false-discovery rate at $\alpha = 0.05$).

Figure S8 shows the decoding profile of classifiers trained on power, phase and voltage extracted from EOG data. The only consistent significant differences were between voltage and the EOG analogue of voltage.

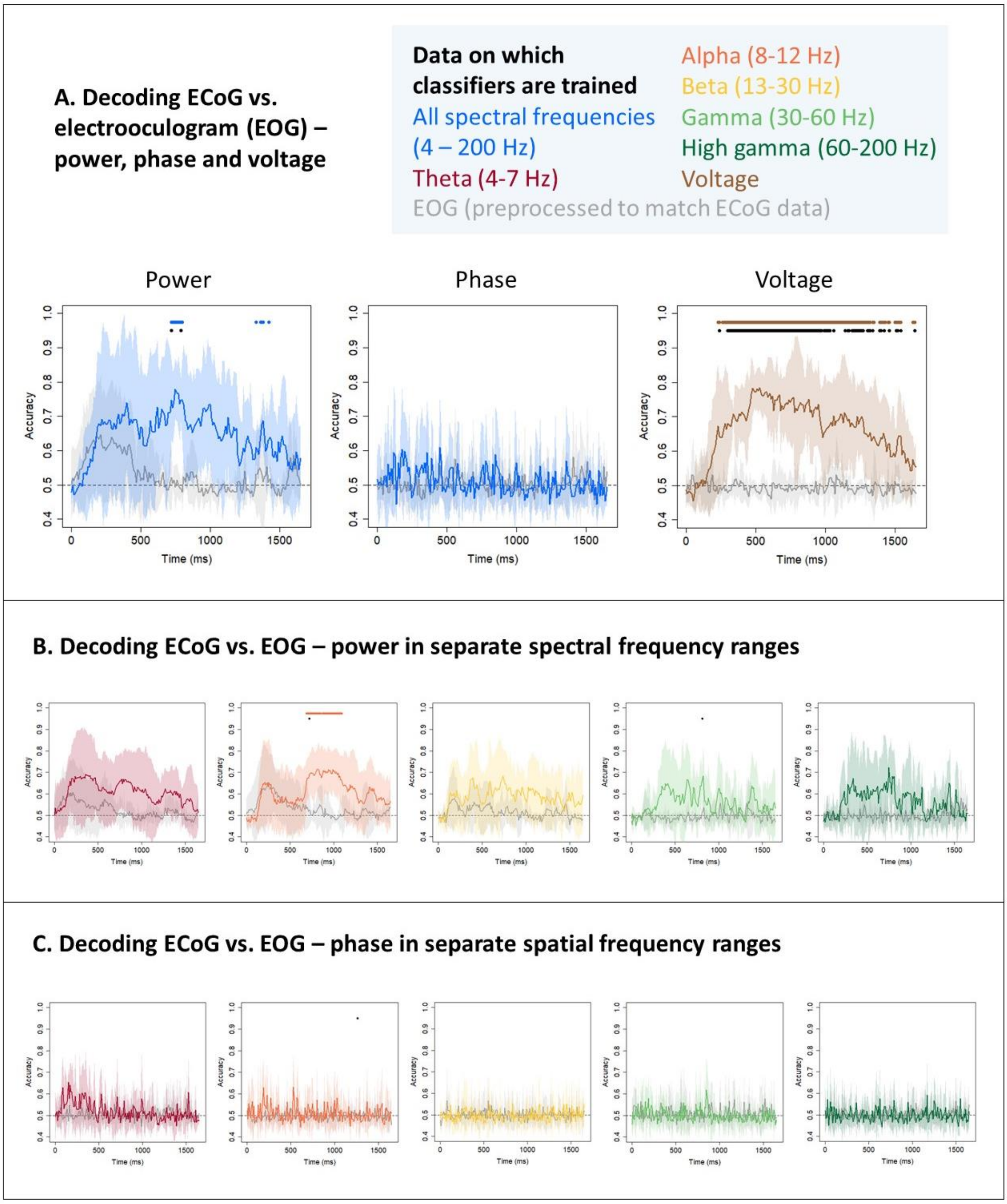


Figure S8: Decoding electrooculogram (EOG) data. (A) Mean and 95% confidence interval of the hold-out accuracy for classifiers trained on power or phase frequency features for all frequencies between 4 and 200 Hz (blue) or on voltage features (brown), compared to classifiers trained on frequency features or on voltage features extracted from EOG data (grey). Coloured dots indicate a significant difference between classifier accuracy and chance (0.5, one-sample t-test with probabilities adjusted to control the false-discovery rate at $\alpha = 0.05$). Black dots indicate a significant difference between accuracies at a given timepoint (paired t-tests with probabilities adjusted to control the false-discovery rate at $\alpha = 0.05$). (B) Mean and 95% confidence interval of the hold-out accuracy for classifiers trained on frequency features composed only of power values extracted from ECoG data within a given range - theta (4 – 7 Hz, red), alpha (8 – 12 Hz, orange), beta (13 – 30 Hz, yellow), gamma (30 – 60 Hz, light green) and high gamma (60 – 200 Hz, dark green) - compared to classifiers trained on frequency features composed of power or phase values extracted from EOG data within that same range (grey). Coloured dots indicate a significant difference between classifier accuracy and chance; black dots indicate a significant difference between accuracies. (C) Mean and 95% confidence interval of the hold-out accuracy for classifiers trained on frequency features composed only of phase values extracted from ECoG data within a given range - theta (4 – 7 Hz, red), alpha (8 – 12 Hz, orange), beta (13 – 30 Hz, yellow), gamma (30 – 60 Hz, light green) and high gamma (60 – 200 Hz, dark green) - compared to classifiers trained on frequency features composed of power or phase values extracted from EOG data within that same range (grey). Coloured dots indicate a significant difference between classifier accuracy and chance; black dots indicate a significant difference between accuracies.

### 4. Discussion

We investigated two aspects of the preprocessing pipeline: the effect of removing noise and saccadic activity.

First we found that heavier cleaning hardly affected decoding with time-frequency power, subtly affected decoding with time-frequency phase, and had a significant effect on decoding with voltage. The absence of an influence of noise on decoding with time-frequency power was important to establish given that artefacts in the data could be "smeared" across (and therefore obscure) true signal during wavelet convolution. One possible reason for this result that we used LASSO (L1) regularisation (Tibshirani, 1996), which selects a sparse subset of the features offered to the classifier. Features obscured by artefacts may be removed by the classifier and decoding performance will be good as long as enough unobscured features remain. Regularisation methods that assume that the code is dense, such as ridge (L2) regularisation (Hoerl & Kennard, 1970), are forced to place weights on all features and so the presence of artefacts may be deleterious to decoding when a different type of classifier is used.

The significant difference for voltage appeared after filtering: before filtering, the decoding profile resembled that obtained by (Rogers et al., 2021). Three filters were applied to the data – CleanLine filtering (Mitra & Bokil, 2007) for line noise at 60 Hz and the harmonics, high-pass filtering at 0.5 Hz, and low-pass filtering at 300 Hz. For 9 patients whose data were recorded at 1000 Hz, the low-pass filter was imposed by the recording equipment – since this was the same for (Rogers et al., 2021), who used no low-pass filter, the low-pass filter is unlikely to be responsible for the decrease in accuracy. Line noise is constant over time and so does not differ between living and nonliving trials. Therefore, the culprit is likely to be the high-pass filter – filtering very low-frequency activity out may change the shape of the voltage deflection over time and reduce its correlation with animacy.

We found that there were few significant differences between the decoding profiles of power and phase extracted from ECoG data and the decoding profiles of power and phase extracted from EOG channels. It should be noted that it is difficult to draw firm conclusions from this analysis because the sample size was small – only 5 of our 19 patients had EOG data. A larger sample size may reinforce the conclusion that there is no significant difference. Alternatively, on visual inspection, the profiles of EOG decoding and ECoG decoding appear to differ and this difference may solidify with a larger number of participants. Until we have clarified the extent to which eye movements may contain information about semantic category, we should take care to correct for eye movements as best we can. Future ECoG data collection projects should take care to record EOG data so that they can be subtracted from the ECoG signal.

To summarise, preprocessing aims to eliminate the possibility of false-positive results (for example, significant decoding driven by a difference in eye movement between categories) and false-negative results (for example, artefacts destroying signal during time-frequency decomposition). We found evidence that most stages of preprocessing make little, if any, difference to time-frequency power and phase decoding with logistic regression with L1 (LASSO) regularisation. Filtering can remove signal as well as noise from voltage data; future work should avoid, or exercise caution when, applying heavy preprocessing to voltage data (Delorme, 2023). Evidence for significant ECoG decoding above that which can be achieved with EOG is inconclusive; future work should seek to clarify the extent to which EOG can cause false positive decoding.

### References


de Cheveigné, A. (2023). Is *EEG is better left alone?* https://doi.org/10.1101/2023.06.19.545602

Delorme, A. (2023). EEG is better left alone. *Scientific Reports*, *13*(1), 2372. https://doi.org/10.1038/s41598-023-27528-0

Hoerl, A. E., & Kennard, R. W. (1970). *Ridge Regression: Biased Estimation for Nonorthogonal Problems*.

Humphreys, G. W., & Forde, E. M. E. (2001). Hierarchies, similarity, and interactivity in object recognition: “Category-specific” neuropsychological deficits. *Behavioral and Brain Sciences*, *24*(3), 453–476. https://doi.org/10.1017/S0140525X01004150

Mitra, P., & Bokil, H. (2007). *Observed Brain Dynamics*. Oxford University Press. https://doi.org/10.1093/acprof:oso/9780195178081.001.0001

Rogers, T. T., Cox, C. R., Lu, Q., Shimotake, A., Kikuchi, T., Kunieda, T., Miyamoto, S., Takahashi, R., Ikeda, A., Matsumoto, R., & Lambon Ralph, M. A. (2021). Evidence for a deep, distributed and dynamic code for animacy in human ventral anterior temporal cortex. *eLife*, *10*, e66276. https://doi.org/10.7554/eLife.66276

Tibshirani, R. (1996). Regression Shrinkage and Selection Via the Lasso. *Journal of the Royal Statistical Society: Series B (Methodological)*, *58*(1), 267–288. https://doi.org/10.1111/j.2517-6161.1996.tb02080.x
